# Of mice and men: Dendritic architecture differentiates human from mice neuronal networks

**DOI:** 10.1101/2023.09.11.557170

**Authors:** Lida Kanari, Ying Shi, Alexis Arnaudon, Natalí Barros-Zulaica, Ruth Benavides-Piccione, Jay S. Coggan, Javier DeFelipe, Kathryn Hess, Huib D. Mansvelder, Eline J. Mertens, Julie Meystre, Rodrigo de Campos Perin, Maurizio Pezzoli, Roy Thomas Daniel, Ron Stoop, Idan Segev, Henry Markram, Christiaan P.J. de Kock

## Abstract

The organizational principles that distinguish the human brain from other species have been a long-standing enigma in neuroscience. Focusing on the uniquely evolved human cortical layers 2 and 3, we computationally reconstruct the cortical architecture for mice and humans. We show that human pyramidal cells form highly complex networks, demonstrated by the increased number and simplex dimension compared to mice. This is surprising because human pyramidal cells are much sparser in the cortex. We show that the number and size of neurons fail to account for this increased network complexity, suggesting that another morphological property is a key determinant of network connectivity. Topological comparison of dendritic structure reveals much higher perisomatic (basal and oblique) branching density in human pyramidal cells. Using topological tools we quantitatively show that this neuronal structural property directly impacts network complexity, including the formation of a rich subnetwork structure. We conclude that greater dendritic complexity, a defining attribute of human L2 and 3 neurons, may provide the human cortex with enhanced computational capacity and cognitive flexibility.

**Graphical abstract:** 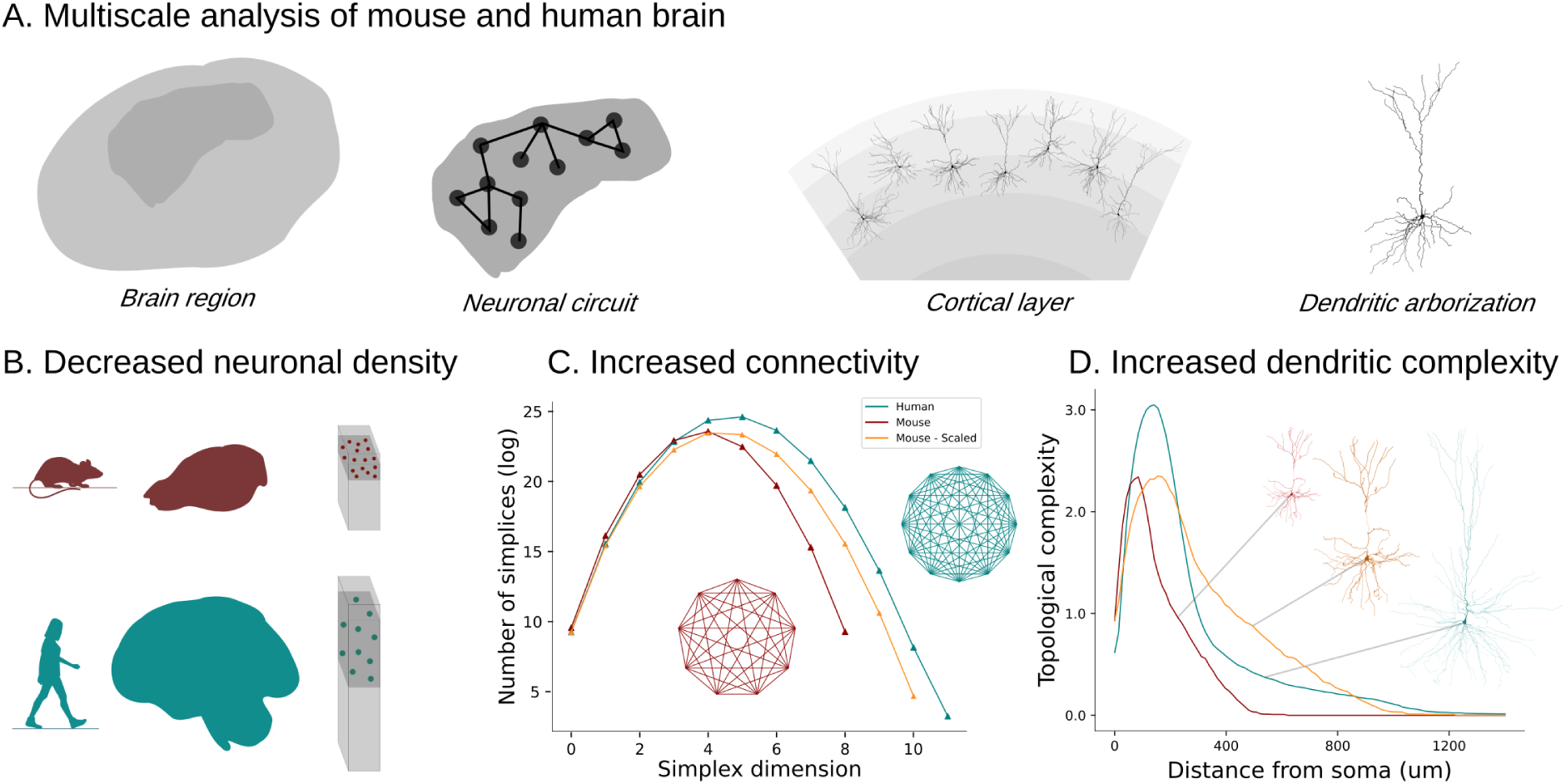

A. A multiscale analysis was performed to compare the mouse and human brains: from the anatomical properties of brain regions to the morphological details of single neurons. B. Human circuits are larger than mice in terms of size and number of neurons, but present decreased neuron density, resulting in increased distances between neurons, particularly among pyramidal cells. C. Greater network complexity emerges within the human brain. Network complexity is defined by larger groups of neurons forming complex interconnections throughout the network. D. The topological analysis of layer 2/3 pyramidal cells in the temporal cortex reveals an intriguing difference: human neurons exhibit a significantly larger number of dendritic branches, especially near the cell body compared to mice. This phenomenon is termed ”higher topological complexity” in dendrites. Our findings suggest that dendritic complexity wields a more substantial influence on network complexity than neuron density.

## Introduction

The question of how the brain contributes to human cognition has been a topic of debate and discussion since ancient times. The shift from Aristotle’s belief that intellect resided in the heart to Gallen’s assertion of the brain’s significance in ancient Rome marked a pivotal moment in this ongoing discourse. Humans have long been intrigued by the reasons behind their relative sophistication compared to other animals (1), sparking deep curiosity about the unique capabilities of the human brain. Initially, it was postulated that human cognitive prowess was linked to the sheer magnitude of our brains (1), stemming from the notion that intelligence is directly correlated with brain size. However, research has failed to identify similar abilities in other large-brained animals, such as elephants (2) and cetaceans (3). Furthermore, subsequent studies (4) systematically analyzing the ratio of brain to body mass refuted these theories, demonstrating that human brain size is not distinctive compared to other animals. Nevertheless, the conviction that the human brain is exceptional among mammalian brains persists, as indicated by the numerous studies that investigate possible correlations of human intelligence with larger cell counts (5), cortical thickness (6; 7), increased cortical folding (8) or dendritic size (9). However, despite extensive endeavors to unravel its mysteries, numerous aspects of our unique characteristics remain elusive. While there are several other factors at play in defining human intelligence, in this study we demonstrate that the shapes of dendrites are an important indicator of network complexity that cannot be disregarded in our quest to identify what makes us human.

Santiago Ramon y Cajal sparked the intriguing inquiry about the distinctive structural characteristics in human neurons that contribute to their optimal functioning compared to neurons in other species (10). Numerous studies have examined the unique functional, molecular, and structural features of human neurons (11; 12; 13). However, it is only recently that significant progress has been made in addressing the knowledge gap concerning distinctive features of human neurons by integrating multimodal datasets that encompass single neuron morphologies, neurochemistry, electrophysiology, and transcriptomics (14; 15; 16; 17; 18). Several laboratories (12; 13; 19; 15) have contributed valuable insights into the structural properties of human neurons by generating morphological reconstructions. In parallel, detailed electron microscopy (EM) reconstructions of human brain tissue (20; 21; 22; 23; 24) have provided important information regarding the composition and connectivity properties of human neurons. These recent advancements in data generation enable a more comprehensive exploration of the intricate structural properties of human neurons and their potential implications for neural function.

Previous analysis of the anatomical properties of the cortex, based on dense tissue reconstructions (22) and detailed anatomical studies (11; 25) indicated a significantly lower cell density in the human cortex, leading to greater inter-neuron distances. In addition, the percentage of interneurons is increased substantially in the human temporal cortex compared to the rodent (11; 25; 22). Putting together the anatomical properties of the mouse and human cortex (11; 25; 22) and the experimental reconstructions of pyramidal cells in layers 2 and 3 of mouse and human cortex, we generated representative networks of both species. Our investigation revealed that the combination of lower neuronal density and the unique morphological properties of human neurons resulted in a considerable increase in the number of directed simplices (26; 27) within the subnetworks of human pyramidal cells, suggesting the presence of abundant, strongly connected subnetworks within the human cerebral cortex. We therefore embarked on a thorough investigation to unravel the underlying factors and discern the distinctive features of human pyramidal cells that result in these significant differences in network complexity between species.

Individual neurons must possess greater dendritic length to maintain the neuropil density within the cortical tissue, despite the greater inter-neuronal distances. This hypothesis was initially proposed based on old histological studies (by Franz Nissl and Constantin Von Economo), suggesting that the wider separation of neurons in humans, compared to other species, could indicate a higher level of refinement in the connections between neurons (28). Through the morphological analysis of neurons, we corroborated previous findings (12; 19; 14) indicating that human neurons exhibit greater total length and extend further from the soma. However, we established that size scaling (14), whether uniform or not, fails to adequately account for the observed increased connectivity in the subnetworks of human pyramidal cells. Our topological analysis (29) unveiled a distinctive pattern of high branching density surrounding the soma in human neurons, a trait absent in the mouse cortical dendrites. This property generalizes across other cortical layers (e.g., layer 5, temporal association cortex) and brain regions (e.g., CA1 hippocampus, pyramidal cell layer), implying that this topological difference may be a universal feature distinguishing human pyramidal cell morphologies from those of mice. This distinctive property proved pivotal in elucidating the observed network disparities. The distribution of additional dendritic branches near the soma confers the advantageous ability to fill the inter-neuronal space while preserving the connection probability between neurons (30) and forming strongly connected cortical subnetworks of pyramidal cells. In addition, the abundance of these strongly connected pyramidal cell cliques necessitates the over-expression of inhibitory neurons (22), which regulate cortical excitation through strongly connected interneuron subnetworks. The findings of our study suggest that a fundamental geometric principle underlies the observed variations in neuronal characteristics, which has significant implications for the structural organization of human networks. Specifically, human excitatory networks prefer a greater complexity of individual cells, as opposed to the larger neuron density observed in mouse networks. This result does not generalize to the subnetwork of interneurons, which are individually less complex, in terms of morphological branching (31; 32), but more abundant in humans, raising further questions about the balance between excitation and inhibition in different species. Our findings shed light on mechanisms underlying the exceptional cognitive abilities exhibited by humans and highlight the significance of considering the interplay between neuronal complexity and network organization in understanding brain function.

## Results

### Human cortex has lower neuronal density and larger inter-neuron distances

Contrary to previous assumptions, the size of the brain is not a proxy for the number of neurons it contains (5). In particular, despite the increase in cell counts observed in the human brain (11), the lower neuron density in the human temporal cortex (≈ 25, 700*/mm*^3^) compared to the mouse temporal cortex (≈ 137, 600*/mm*^3^) challenges this notion (ure 1B). Another important difference is the significantly higher proportion of interneurons (Figure 1C), i.e. inhibitory neurons with local dendrites, comprising 30% of human neurons compared to a mere 12% in the mouse (28; 22).

**Figure 1:**
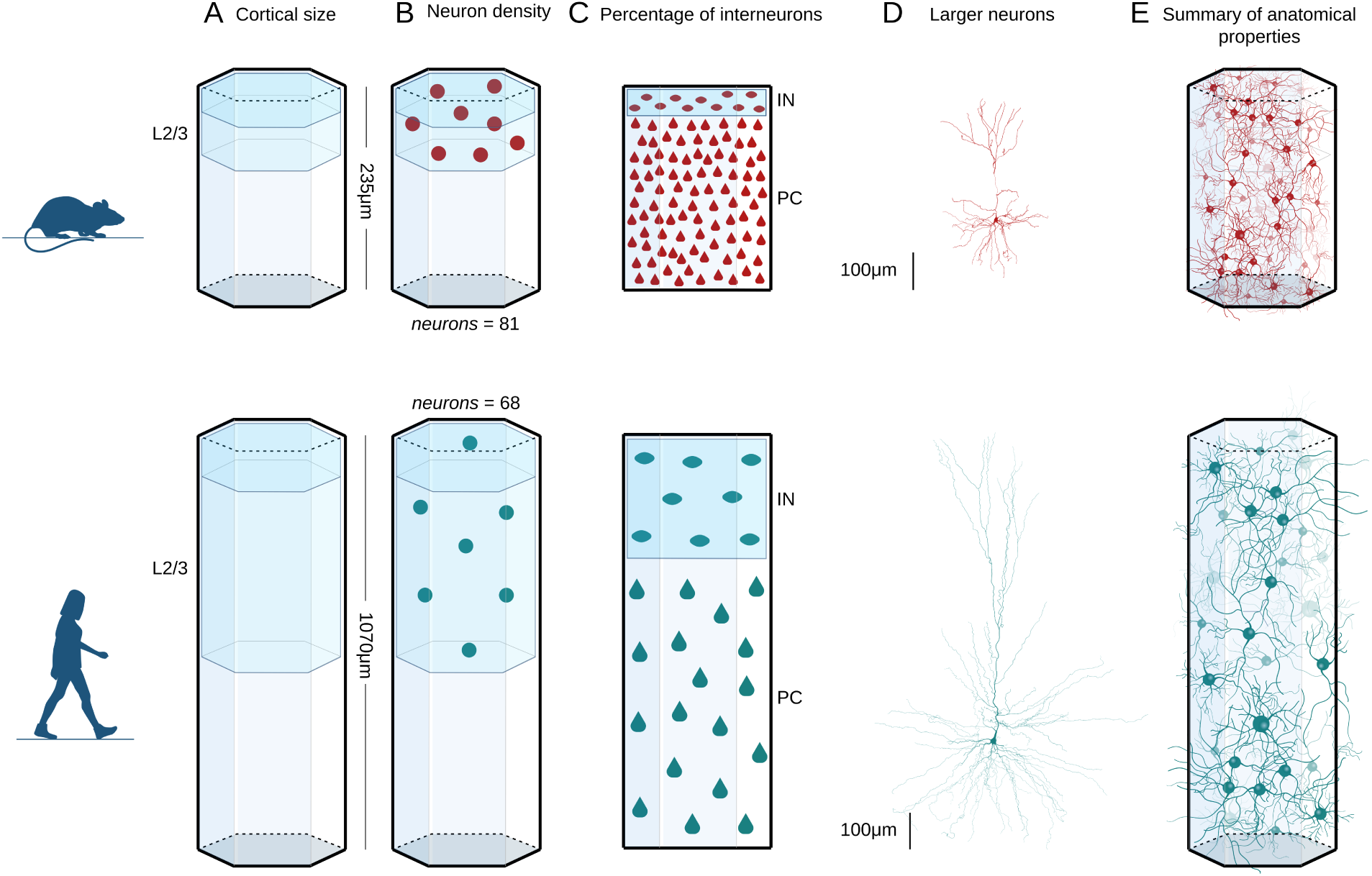
Comparative analysis of network architecture. A. Lower cortical thickness in mouse (top) compared to human (bottom), results in longer cortical layers 2 and 3 (L2/3) in human. B. Lower neuron density in the human cortex leads to increased inter-neuron distances and a lower number of neurons in humans. C. The percentage of interneurons (IN) compared to the pyramidal cells (PC) increases in the human cortex, indicating a significant network difference between species. D. Human neurons are larger with more extended dendrites, as expected by the larger cortical thickness. E. The total dendritic lengths in the cortex are similar between mice and humans balanced by the lower neuron density but larger neurons in humans.

To investigate the effect of lower neuron density on the spatial distribution of neurons, we computed the distance between the cells, assuming a homogeneous distribution of neurons within the cortical layers 2 and 3. The closest neighbor distance between human neurons is ≈ 19*µm*, which is nearly twice that of mice at ≈ 11*µm* (see Methods: Computation of inter-neuron distances). This difference is more striking when computed for pyramidal cells (closest neighbor pyramidal cell distance is ≈ 21.3*µm* in human versus ≈ 11.5*µm* in mouse), as opposed to the distance between interneurons, which is ≈ 28.5*µm* in human versus ≈ 22*µm* in mouse, due to the higher proportion of interneurons in the human temporal cortex (Figure 1C).

We then computed the expected neuropil densities, i.e. the total length of dendritic branches per volume, within the tissue of the two species. For this experiment, we used the number of synapses (11.8 × 10^8^*/mm*^3^ in humans versus 26.56 × 10^8^*/mm*^3^ in mice) within a volume of 1*mm*^3^ of the cortex (11) and the reported synaptic densities (22), 0.88*/µm* in human versus 2.15*/µm* in mouse). This analysis revealed that the total length of dendrites per cubic millimeter is comparable between the two species, with human dendrites measuring ≈ 1, 340*m/mm*^3^ compared to ≈ 1, 240*m/mm*^3^ in mice (Figure 1E). Despite the total dendritic length in the neuropil being comparable between the two species, the total dendritic length per neuron is increased (measure a total dendritic length of 11600 ± 5500 in human versus 5500 ± 2200 in mouse) as observed by the larger size of human arborizations (Figure 1D) (12; 13), due to the significant differences in neuron density (25.7*K* neurons */mm*^3^ in humans versus 137.6*K* neurons */mm*^3^ in mouse).

### Human pyramidal cells generate strongly connected subnetworks

To study how the anatomical differences between the two species contribute to the network architecture, we generated mouse and human networks of the same radius 476*µm* and studied their structural connectivity. The cortical circuits consist of layers 2 and 3 that differ in thickness between the two species (1070*µm* in humans versus 235*µm* in mice, Figure 1A).

Using available data on neuronal densities (11) and the ratio of excitatory to inhibitory neurons (22), along with morphological reconstructions for four different cell types (human pyramidal cells, mouse pyramidal cells, human interneurons, and mouse interneurons, Table S1), we homogeneously distributed neurons and populated the networks with morphological reconstructions. We collected morphologies from diverse sources (12; 13; 19; 15) and ensured the coherence of the datasets by comparing their variation against the within-group variation. Neurons were selected exclusively from analogous brain regions and within comparable age ranges. The total number of neurons in the respective networks is 4503 interneurons and 10621 pyramidal cells in the human and 2724 interneurons and 14377 pyramidal cells in the mouse. The connectivity of the human and mouse networks was computed from the touches of axons to dendrites, i.e., the appositions, determined by branches coming closer than 2*µm*. There is an ongoing debate on the appropriate selection of synapses from the pool of potential synapses (33; 34; 35). It is important to note that the connectivity was determined through a random selection of appositions until a 4% network density was reached in both networks. Due to the random pruning of connections, no direct comparison can be made to the connectivity measurements observed in actual biological circuits. However, the appositions are highly informative on the potential pool of synapses for the respective biological networks.

First, we analyzed the spatial distribution of neurons in the two species (Figure 2A-B). Neurons were distributed homogeneously according to the respective neuron densities. The number of neurons found at different distances (0 − 1000*µm*) from a central neuron was computed (Figure 2B) and demonstrates that fewer neighbors are localized around a human neuron, as expected due to the lower neuronal density in the human cortex. The network density, i.e., the ratio between the actual connections in the network over the maximum number of possible connections (equal to the square of number of nodes), of the human pyramidal cells subnetwork (Figure 2C) is similar to the respective subnetwork in mice. On the other hand, the respective network density of the human interneuron subnetwork is higher, due to the higher percentage of interneurons in the human neocortex (30% in humans versus 12% in mice). In addition, the in-degree distribution(Figure 2E), with a mean and standard deviation of 538 ± 250, in the subnetwork of human pyramidal cells is significantly smaller to the mouse subnetwork (Figure 2E), with a mean and standard deviation of 712 ± 239 (*p* ≈ 0), and it is significantly increased in the subnetwork of human interneurons (Figure 2F, 95 ± 69 in human versus 28 ± 13 in mouse, *p* ≈ 0). The details of the mouse and human network properties are summarized in Table S2.

**Figure 2:**
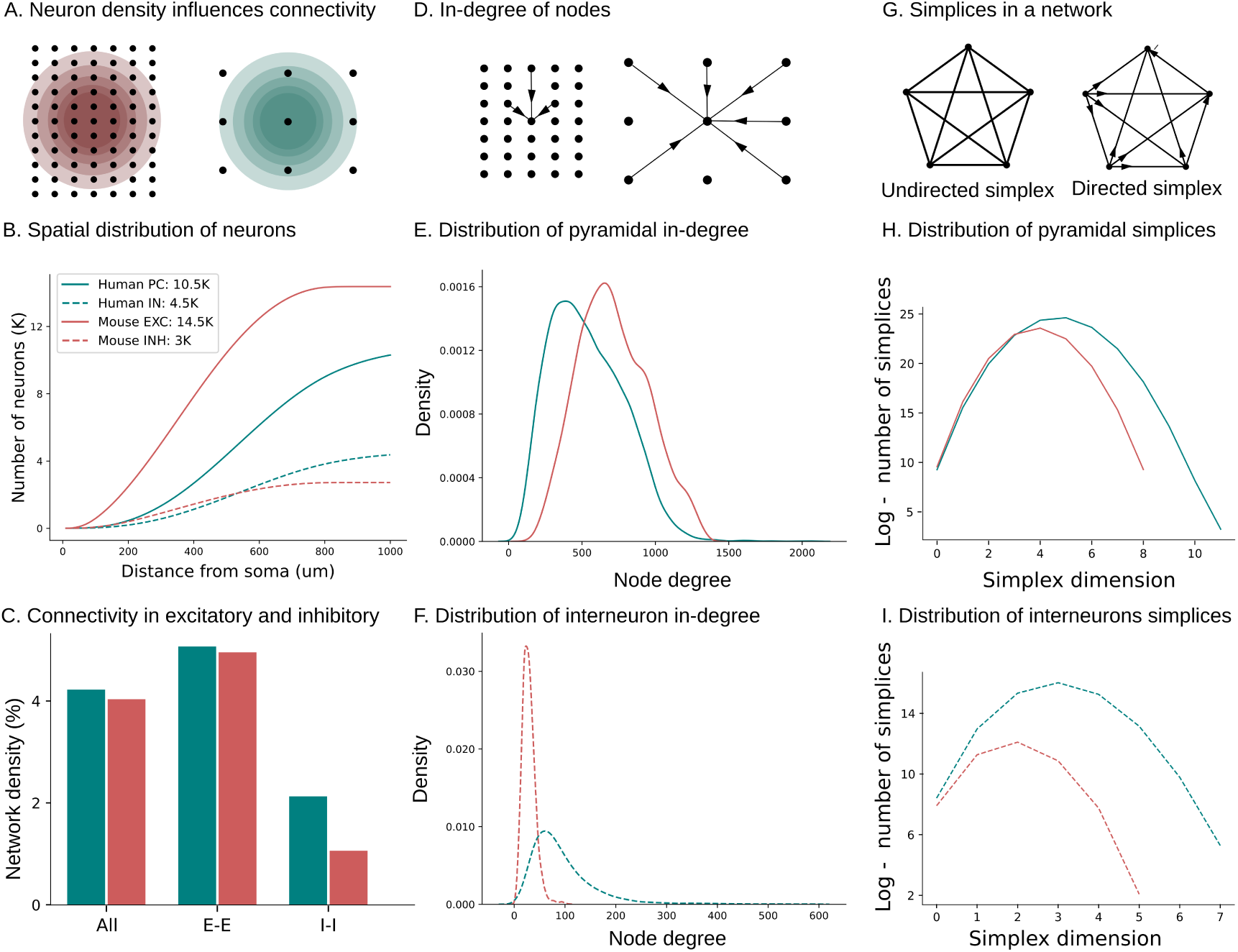
Human pyramidal cells generate strongly connected subnetworks. A. Grid illustrating differences in neuron density in mice (red) and humans (teal). Concentric spheres surrounding a central neuron portray the cell counts presented in (B). B. Number of neighbors. A cortical layer 2/3 of dimensions 1*mm* x 1*mm* was simulated for the respective cortical thickness of mouse (235*µm*) and human (1070*µm*) and the respective neuron counts for mouse (28*K*) and human (18*K*). The number of neurons at different distances (0 − 1000*µm*) from a central neuron was computed. Insert reports of total cell counts. Cell counts do not linearly correspond to cell density due to the differences in cortical thickness, which results in a larger cortical volume in the human cortex. C. Network density (number of edges over maximum possible number of edges) for all neurons, and subnetworks of pyramidal cells and interneurons. D. Schematic of node in-degree for mouse and human networks. E. Distribution of in-degrees for and mouse (red) and human (teal) pyramidal cells subnetworks. F. Distribution of in-degrees for mouse (red) and human (teal) interneuron subnetworks. G. Example of a five-simplex in undirected and directed graphs. H. Distribution of simplices for subnetworks of pyramidal cells. Log scale is used for the number of simplices to depict the orders of magnitude higher simplex counts in humans. I. Distribution of simplices for subnetwork of interneurons.

The complexity of the networks was assessed by computing the distribution of directed simplices (27). Simplices provide a comprehensive representation of fully connected subgraphs within the networks (Figure S1), combining information about the density and the degree distribution of the graphs and represent network hubs that are related to modulation of cortical dynamics (36; 37). Furthermore, it is worth noting that higher-dimensional simplices are associated with increased robustness in networks (38), since information can be efficiently transmitted in a specific direction, as all edges within the simplices are aligned to transmit information in the same direction. Surprisingly, despite the lower neuronal density observed in the human layer 2/3 cortex, the subnetwork of pyramidal cells exhibited a complex connectivity pattern, leading to the formation of higher-dimensional simplices. Specifically, in the human network, we observed simplices of dimension 12, whereas, in the mouse network, the dimension was limited to 9 (Figure 2H). The number of simplices in the human network was approximately three times larger compared to the mouse network. These findings suggest that the human cortical network displays a greater degree of complexity than its mouse counterpart, despite the lower neuronal density. These results generalize to the network of interneurons (Figure 2I), due to the larger inhibition/excitation ratio in humans. Interneuron subnetworks exhibit higher dimensional simplices (8 in humans versus 6 in mice) and even larger simplex counts (Figure 2I). Our results suggest that human cortical networks consist of an abundance of spatially sparse high-dimensional directed cliques of pyramidal cells which are controlled by a dense subnetwork of interneurons.

### Simple dendritic scaling cannot explain species-specific phenotypes

We then investigated which morphological features explain the significant increase in neuronal connectivity in human networks. We evaluated whether a simple scaling law can determine the relationship between the dendrites of the two species. Based on the observed variations in neuron size (14) and the conserved molecular cell types (39; 15; 16) across different species, the existence of a simple scaling law governing morphological cell types has been hypothesized. Due to the observed differences in body size, which results in larger human brains and increased cortical thickness (Figure 3A), it is assumed that the larger cell counts and the increased dendritic lengths can explain the enhanced cognitive abilities of humans (40; 9; 41).

**Figure 3:**
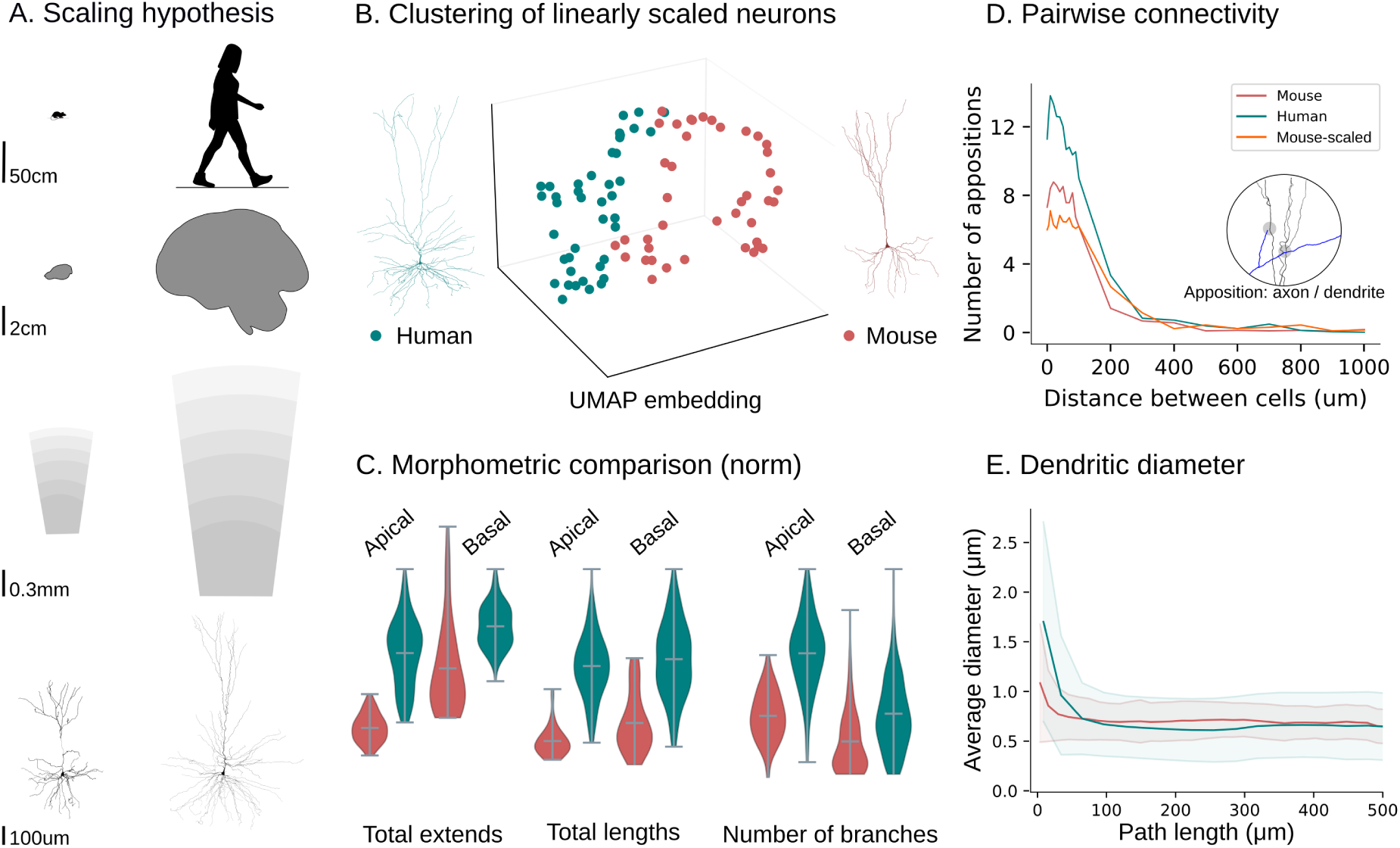
Simple scaling cannot explain species-specific phenotypes. A. Schematic representation of the scaling hypothesis: the dendritic size scales with the brain size, which in turn scales with the body size. B. The UMAP embedding of the topological representations (normalized to the same total size) of mouse (red) and human (teal) neurons shows a distinction between the two species. C. Comparison of pyramidal cell morphometrics (normalized by maximum value in human, separated in apical and basal dendrites) show that morphological features do not scale uniformly. D. Pairwise number of appositions at different inter-neuron distances for pairs of mouse (red), human (teal), and scaled morphology of mouse (orange) neurons. E. Average dendritic thickness (diameters) are comparable between mice and humans at path distances above 150*µm*. However, close to the soma diameters are almost double in human pyramidal cells.

Morphologies collected from diverse sources (12; 13; 19; 15) were analyzed in terms of morphometric (42) and topological (29) properties. The datasets employed for our analysis consisted of mouse and human pyramidal cells (from the temporal cortex (layers 2, 3 and 5) and hippocampus and interneurons from the layers 2 and 3 neocortex (see SI: Datasets, Table S1). A comprehensive analysis to check biases in the data was also performed (Figures S2, S3). There was no bias in terms of sex, condition and age (Figure S2). Interestingly, neurons from the same patients showed lower variance (Figure S3) indicating that individuals may posses unique morphological traits, a perspective that could lead to personalized medicine in the future. Initially, we focused on pyramidal cells from layers 2 and 3. The clustering of human neurons and scaled mice neurons to compensate for the difference in cortical thickness between the two species (Figure 3B) demonstrates that cortical thickness alone cannot explain the observed morphological differences.

A comprehensive comparison of morphological characteristics between the pyramidal cells of cortical layers 2 and 3 of the two species revealed profound differences between human and mouse neurons (Figure 3C) and confirmed findings reported in previous studies (13; 19). In the following measurements, we report mean, standard deviation, and p-values based on a KS-statistical test between the two distributions. Human pyramidal cells are larger, evidenced by their increased total lengths both in apical (5958 ± 2636*µm* in human apical dendrites, versus 2523 ± 1021*µm* in mice, *p <* 10*^−^*^15^) and basal dendrites (5644 ± 3921*µm* in human basal dendrites, versus 2952±1525*µm* in mice, *p <* 10*^−^*^5^). In addition, human neurons extend to larger distances (692 ± 272*µm*) compared to mice (349 ± 96*µm*, *p <* 10*^−^*^11^) due to the increased thickness of the human cortex (2.2 times larger) and particularly layers 2/3 (4.5 times larger in human). Additionally, human apical dendrites possess more branches (54±21 in humans, versus 36 ± 11 in mice, *p <* 10*^−^*^5^) and human basal dendrites have also increased number of branches (61 ± 34 in humans, versus 46 ± 19 in mice, *p* = 0.006). The detailed morphological analysis is summarized in Tables S3-S6. Another significant difference is the increase in dendritic thickness close to the soma for human dendrites (Figure 3E) despite the comparable mean thickness values between the two species (0.34 ± 0.11 in human apical dendrites compared to 0.38 ± 0.06 in mouse, *p <* 10*^−^*^5^, and 0.32 ± 0.11 in human basal dendrites compared to 0.34 ± 0.04 in mouse, *p* = 0.00025). In this case, the statistical differences do not come from the mean values, but from the differences in the distributions as demonstrated by the maximum diameters (0.68 in human apical dendrites compared to 0.56 in mouse, 0.65 in human basal dendrites compared to 0.45 in mouse) which typically occur near the soma.

A computation of pairwise connectivity at varying distances between pre-synaptic and post-synaptic cells (Figure 3D) shows a higher number of appositions, i.e., touches that can potentially become synapses, in human cells than in mouse cells. Surprisingly, when a similar analysis was conducted on mouse cells scaled to twice their original extents, fewer appositions were observed, an effect that can be attributed to the increased space between branches. Therefore, the increased total lengths of dendrites cannot sufficiently explain the observed connectivity differences. We hypothesize that the spatial arrangement of branches and their organization around the soma are important determinants of neuronal connectivity.

### Topological profiles of layer 2 and 3 pyramidal cells show species-specific morphological traits

To identify the precise morphological features that are unique in each species, we studied the topological properties of neurons. The topological morphology descriptor, TMD (29), which encodes the topology of branches at different path distances from the soma, was used to study how branches are distributed in the two species. The extracted topological barcodes of the human (see Figure S4) and mouse (see Figure S5) pyramidal cells unveiled a fundamental difference in their branching patterns (Figure 4D-E). Specifically, collateral dendritic branches of human pyramidal cells (Figure 4A-B) start closer to the soma (200 − 400*µm*, Figure 4F) but extend to larger radial distances, thus conferring a distinct topological profile of longer branches close to the soma in human neurons. In the vicinity of the soma human dendrites of pyramidal cells exhibit a higher density of branches (Figure S 6D), which were also found to be longer than their mouse counterparts (Figure S 6E).

**Figure 4:**
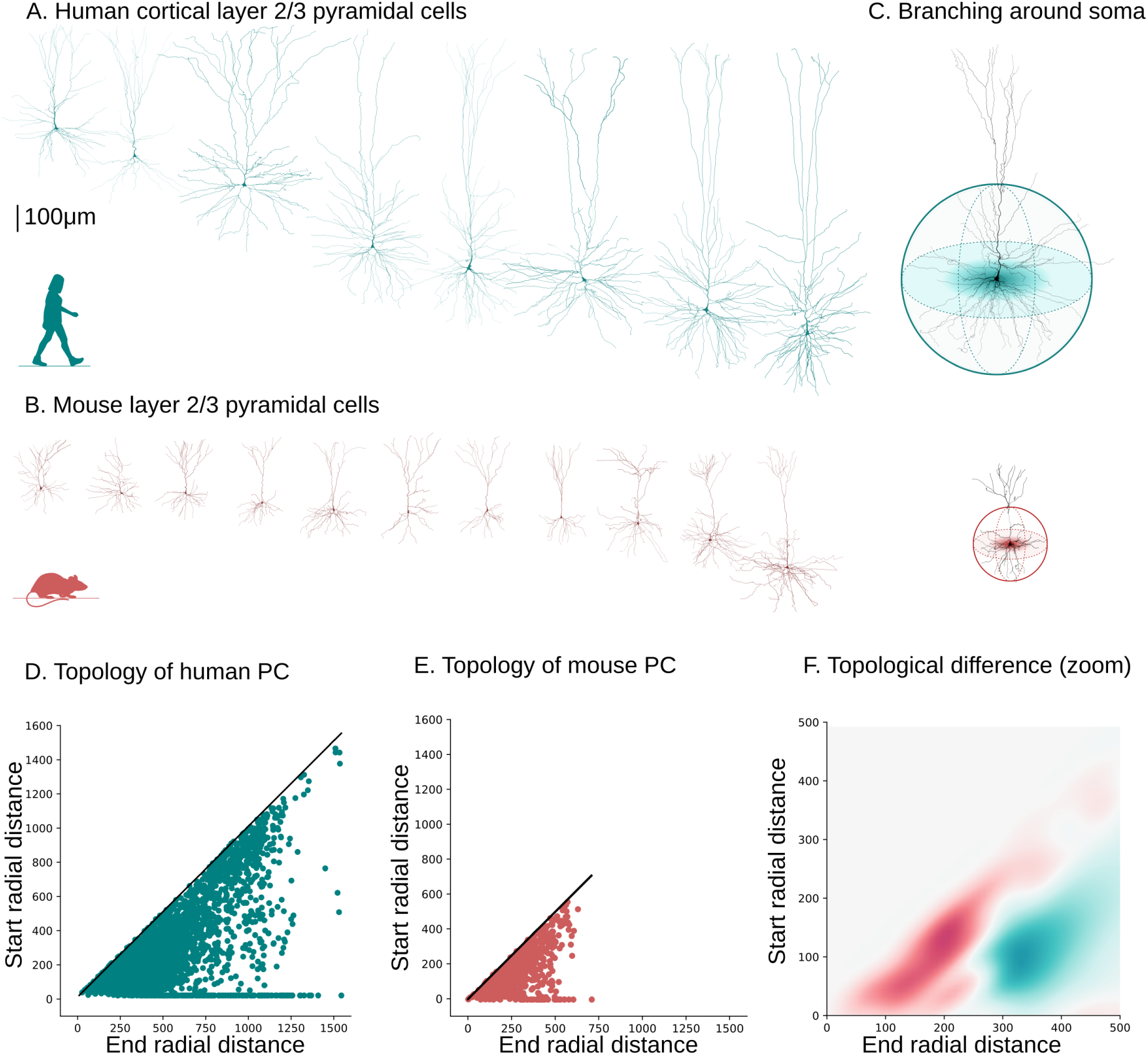
Comparative topological analysis of layer 2 and 3 cortical pyramidal cells. Exemplary reconstructions of layer 2 and 3 pyramidal cells mice (B) and in humans (A). C. Schematic of topological branching around the neuronal soma for mice (bottom) and humans (top). Topological representation of mouse (E) and human (D) neurons and their difference (F) around the soma; zoom in the area of maximum difference at 200 − 500*µm*.

We used an optimization algorithm to approximate the experimental data by minimizing the distances between mouse and human persistence barcodes. However, neither uniform nor non-uniform scaling (Figure S 7A-B, Methods: Scaling optimization) could adequately fit the experimental data. The normalized persistence diagrams were also approximated by Gaussian kernels (three centers, Figure S 7C) in both species. The distinct dendritic density patterns are species-specific, as evidenced by the spatially distinct differences between the approximated Gaussian kernels. Species-specific topology explains why simple scaling laws cannot sufficiently approximate the data. The higher perisomatic branch density is essential to ensure high connectivity between pairs of human neurons, as computed in (30), compensating for the smaller number of neighbors around a human neuron.

Due to the limited sample size of interneuron reconstructions, we focused on basket cells(Figures S8, S9, S10), representing the majority of interneurons in our dataset ≈ 70%. A topological analysis performed on the dendrites of basket cells in layers 2 and 3 for humans and mice (Figure S6) demonstrated that the observed topological differences between the two species persist in the dendrites of interneurons (Figure S6). However, the higher density of branches around the somata is significantly less prominent than pyramidal cells. This distinct topological pattern highlights a clear differentiation between pyramidal cells and interneurons within the cortical circuitry, further emphasizing the diversity of neuronal architecture in the human brain and suggesting that different cell types contribute to the overall network organization in distinct ways.

To clarify the impact of cell density versus neuronal morphology on network properties, we constructed a hybrid network (Figure 5B) for comparison with both mouse (Figure 5A) and human networks (Figure 5C). This hybrid circuit consists of layers 2 and 3 with a human-like thickness of 1070*µm*, human cell densities, and a radius of 476*µm*. The mouse morphologies were scaled to match the total dendritic lengths found in humans, and human axons were attached to these scaled mouse dendrites. As a result, the hybrid circuit resembles a human-like network, with human dendritic morphologies substituted for their mouse-scaled counterparts. The distribution of simplices in the hybrid subnetwork of pyramidal cells closely matches the mouse distribution (see Figure 5D, top), suggesting that dendritic complexity (see Figure 5E, top) is the key determinant contributing to the highly complex human pyramidal cell subnetworks. In contrast, the hybrid subnetwork of interneurons shows a significant increase in the number of simplices compared to the mouse distribution (see Figure 5D, bottom). This finding highlights that given the lower complexity of interneuron dendrites (see Figure 5E, bottom), cell density plays a crucial role in determining network complexity. Another important aspect is axon complexity. As evident by Figure 5F human axons are less complex both in pyramidal cells and interneurons compared to mice. Therefore, we are probably missing a significant portion of human axons due to the difficulty of reconstructing intact axons in human tissue, leading to an underestimation of simplices in the human circuit.

**Figure 5:**
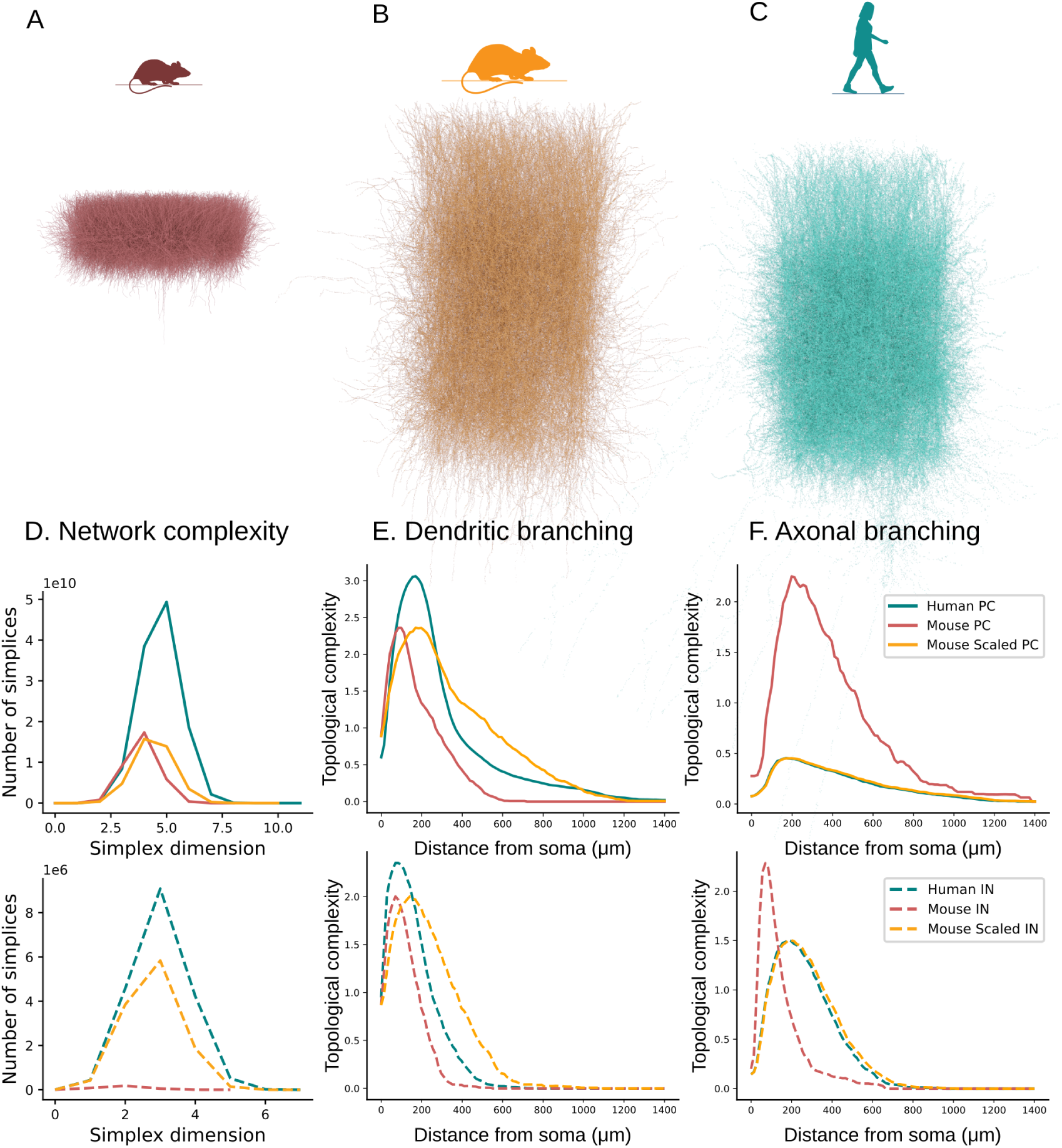
From dendritic complexity to network complexity. Examples of mouse (A - red), hybrid (B - orange), and human (C - teal) layer 2 and 3 circuits with realistic thickness. D. Network complexity based on the distribution of simplices of the pyramidal cell subnetwork (top, solid lines) and the interneuron subnetwork (bottom, dashed lines). E. Dendritic complexity based on the topology of the pyramidal cells (top, solid lines) and the interneurons (bottom, dashed lines). F. Axonal complexity based on the topology of the pyramidal cell (top, solid lines) and the interneurons (bottom, dashed lines).

The topological analysis was generalized to other cortical layers (layer 5) and brain regions (hippocampus) for which data were available. The topological examination of pyramidal cells from the layer 5 temporal cortex revealed the persistence of observed topological differences in deeper cortical layers. The dendritic branches of layer 5 human pyramidal (Figure S11) cells exhibited a higher density in proximity to the soma, with branches commencing earlier and extending further away from the soma than in mice, confirming that the observed topological profiles of human cells extend to neurons beyond layers 2 and 3. Remarkably, these results also generalize to other brain regions. Data obtained from Benavides-Piccione et al (19) in the mouse and human hippocampus (Figure S11) demonstrated a similar pattern. This finding is particularly intriguing as it highlights a distinct single-cell characteristic that serves as a signature feature of human cells, with high branch densities localized at 200 − 500*µm* around neuronal somata.

Finally, we evaluated the effect of the increased number of neurons in the cortex as a determinant of the increased network complexity. We generated random small world networks (Watts-Strogats (43) with probability *p* = 0.7) and random networks (Erdos-Renyi (44)) with the corresponding structural properties of the networks of pyramidal cells (number of nodes and network density) and compared them to the reconstructed networks with similar properties (Figure S12). This analysis indicates that network size (number of nodes and network density) cannot sufficiently explain the observed network complexity differences between human and mouse networks.

## Discussion

Despite the greater inter-neuron distances resulting from lower neuron density in human pyramidal cells (Figure 6A), the distinct topology of their dendritic arbors (Figure 6B) emerges as a mechanism to establish higher-order interactions, as evidenced by the larger number of higher-dimensional simplices in the network (Figure 2, Figure 5). A higher number of simplices indicates strong directionality in the human subnetworks of pyramidal cells, a result supported by recent experimental studies (45) that found lower reciprocal connections in human cortical networks. In addition, interneurons that exhibit higher densities in humans, but similar dendritic characteristics (32) further enhance network complexity. These observations support the balance in inhibition and excitation (Figure S13) as experimentally observed in the innervation of pyramidal cells in the human cortex (22). The intricate morphology of individual neurons significantly influences the properties of the networks they contribute to, providing strong evidence of a direct link between dendritic complexity and the overall structure of neuronal networks.

**Figure 6:**
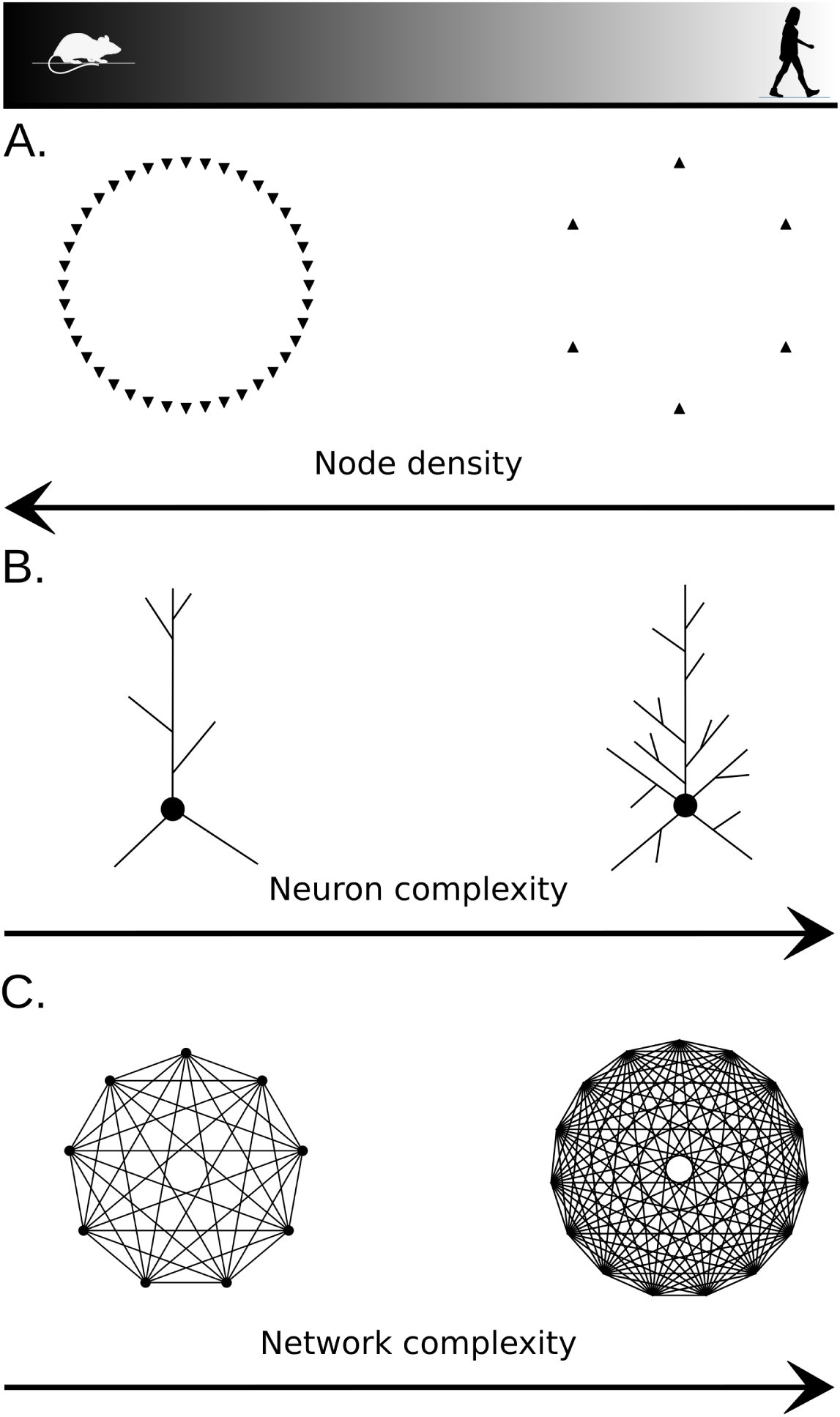
Single-cell complexity influences network complexity. Human networks exhibit a lower density of neurons (A) and therefore larger inter-neuron distances. However, human dendrites (and especially pyramidal cells) are highly complex (B). The combination of complex dendrites in human pyramidal cells and lower neuronal density generates highly complex human subnetworks (C). The effect of increased topological complexity of dendrites is a better indicator of network complexity than node density.

A natural follow-up question is whether the observed structural differences confer a computational advantage to the human brain. Numerous studies have supported the importance of dendritic complexity for computational capacity (46; 47; 40; 48; 49; 50). To address this question, we build upon research in dendritic impact (51) by calculating the anticipated memory capacity for a non-linear branching dendrite and comparing it with its topological complexity (Figure S14). Memory capacity is defined as the number of possible combinations between the inputs of the dendrite and its non-linear components (see Methods: Memory capacity). Topological complexity, in this context, refers to the entropy of bar lengths within the persistence barcode of a neuron (see Methods: Topological analysis). We observed that memory capacity is correlated with topological complexity (Figure S14), as anticipated by the increased number of branches, enabling sampling of a larger combination space for the synaptic inputs. These findings suggest that dendritic complexity is a suitable indicator for predicting the expected memory capacity of individual neurons.

Two primary approaches to achieve higher-order complexity in neuronal networks are commonly proposed: increasing the number of nodes (52) or enhancing the complexity of individual nodes (53; 54). Interestingly, instead of simply adding more nodes, the human pyramidal cells have evolved to prioritize the complexity of individual neurons. This suggests that the human brain has harnessed the potential of single-neuron complexity as a mechanism to create and support complex network connectivity, potentially conferring unique cognitive advantages. This dendritic property combined with the increased perisomatic dendritic thickness could explain electrophysiological findings, such as the lower voltage-gated potassium and HCN conductances identified in (14) and the accelerated signal propagation speed in human pyramidal cells (55). In addition, when considering human cognition, we need to consider differences in the electrophysiological and synaptic properties of the species. Human layer 5 neurons exhibit a unique biophysical composition that sets them apart from other species, as highlighted by Beaulieu-Laroche et al. (14) as well as thicker and longer spines (56). We propose that the branching structure of pyramidal cells, similar to synaptic and electrophysiological properties, does not adhere to traditional allometric rules, emphasizing their importance for future research. Studying the roles of these distinct characteristics and their implications in functional cortical networks is essential for understanding human cognition and pathologies.

Our analysis substantiates a compelling hypothesis that challenges the conventional notion of increasing node density as a single means of optimizing the computational capacity of networks. Instead, we propose that enhancing node complexity, as observed in the distinct morphological features of human neurons, may be an alternative mechanism for improving network performance. The biological mechanism by which increased node complexity enhances the computational capabilities of a network remains an intriguing open question. However, it is not difficult to imagine that the simultaneous greater memory capacity within individual neurons and the substantially higher simplex counts should result in networks with higher computational power. As our understanding of the human neocortex continues to evolve, more detailed models of the human cortex integrating comprehensive information about synaptic, transcriptomic, and morpho-electrical relationships (16; 15), and preferential connectivity rules (17; 22) will create a more accurate understanding of the human brain.

The interplay between interneurons and pyramidal cells, and the functional role of each component within the network, is another important aspect that warrants further investigation. With the discovery of a larger proportion of interneurons in the human brain than in mice, there has been speculation that they may have unforeseen functional significance (22). Our findings suggest that the simpler dendritic shapes of interneurons (32) may necessitate a higher abundance of these cells to maintain the balance between excitation and inhibition within the network, as experimentally observed in (22). However, these results cannot be generalized to all interneuron types, due to the great diversity in interneuron shapes (57; 31) and the specific roles of different cell types in the functionality of human networks (58). Therefore, exploring the connectivity patterns within the whole network of both excitatory and inhibitory cells, rather than focusing solely on their respective subnetworks, would provide valuable insights into the necessity for a greater number of interneurons and their specific contributions to network dynamics (58).

## Methods

### Experimental reconstructions

#### Human Brain Slice Preparation in VU laboratory

All procedures were performed with the approval of the Medical Ethical Committee of the VU medical center (VUmc), and in accordance with Dutch license procedures and the Declaration of Helsinki. All patients provided written informed consent. Tissue was obtained from the Middle temporal gyrus (MTG) during neurosurgical resection of non-pathological cortical tissue, to gain access to deeper-lying pathology. Immediately upon resection, the tissue block was placed into a sealed container filled with ice-cold, artificial cerebral spinal fluid (aCSF) consisting of (in mM): 110 choline chloride, 26 NaHCO3, 10 D-glucose, 11.6 sodium ascorbate, 7 MgCl2, 3.1 sodium pyruvate, 2.5 KCl, 1.25 NaH2PO4, and 0.5 CaCl2. Before use, the aCSF solution was maintained carbogenated with 95% O2, 5% CO2, and the pH was adjusted to 7.3 by the addition of 5M HCl solution.

The transition time between resection in the operation room and arriving at the neurophysiology lab was less than 20 minutes. Immediately upon arrival at the lab, the tissue block was cleaned of residual blood and after identifying the pia-white matter (WM) axis, pia was carefully removed. Subsequently, coronal slices of 350 *µ*m-thick were made using a vibratome (Leica V1200S), in ice-cold aCSF solution. After recovering in a heat bath between 15-30 minutes at 34 °C, the slices were kept for 1 hour at room temperature before recording in aCSF, which contained (in mM): 125 NaCl; 3 KCl; 1.2 NaH2PO4; 1 MgSO4; 2 CaCl2; 26 NaHCO3; 10 D-glucose (300 mOsm), bubbled with carbogen gas (95% O2/5% CO2). Borosilicate glass patch pipettes (Harvard apparatus/Science products GmbH) were filled with intracellular solution, containing (in mM): 115 K-gluconate, 10 HEPES, 4 KCl, 4 Mg-ATP, 10 K-Phosphocreatine, 0.3 GTP, 0.2 EGTA, and biocytin 5 mg/ml, pH adjusted to 7.3 with KOH, osmolarity 295 mOsm/kg). Using this intracellular, the recorded cell passively filled with biocytin during the patch clamp recording, allowing for subsequent staining of the cell.

#### Mouse slice preparation in VU laboratory

All experiments were carried out in accordance with the animal welfare guidelines and approval of the animal ethical committee of the VU Amsterdam, the Netherlands. Adult male and female wild-type C57BL/6J mice (7–15 weeks old) were used and slice preparation procedures were described previously (12; 30). In short, mice were decapitated and their brains were immediately submerged in ice-cold aCSF. Alternatively, mice were anesthetized using Euthasol (4-mg pentobarbital sodium in 0.2*mL* of a 0.9% sodium chloride solution, intraperitoneal injection) and transcardially perfused with 10*mL* of an oxygenated ice-cold choline-based solution. Mice were decapitated and brains submerged in oxygenated choline solution. Coronal sections (350*µm*) including temporal association cortex were cut on a vibratome. After slicing, the same methodology was used for human and mouse samples, including electrophysiological recordings, biocytin filling, histological processing, and reconstruction methods.

#### Immunohistochemistry

Following whole-cell patch-clamp recordings, slices were fixed in 4% paraformaldehyde (in phosphate buffer) for a minimum of two days. Then, biocytin-filled neurons were visualized using the chromogen 3,3-diaminobenzidine (DAB) tetrahydrochloride avidin–biotin–peroxidase method. In addition, a 4,6-diaminidino-2-phenylindole (DAPI) staining was performed. Slices were mounted on slides and embedded in mowiol (Clariant GmbH, Frankfurt am Main, Germany).

#### Morphological reconstruction

All human and mouse neurons underwent critical quality checks including assessment of staining quality and visualization, and occurrence of slicing artifacts (i.e. slicing plane not parallel to apical dendrites). Only neurons that passed quality control were reconstructed with Neurolucida software (Microbrightfield, Williston, VT, USA), using a 100x oil objective. From these 3D reconstructed neurons, morphological parameters could be extracted for further analysis.

#### Human Slice preparation in EPFL laboratory

All procedures were performed with the approval of the Swiss Research Ethical Committee and in accordance with Swiss procedures. All patients provided written informed consent. Tissue was obtained from the idle temporal cortex during neurosurgical resection of non-pathological cortical tissue, to gain access to deeper-lying pathology. Immediately upon resection, the tissue block was placed into a sealed container filled with ice-cold, artificial cerebrospinal fluid (aCSF). During transport, the aCSF solution was maintained carbogenated with 95% O2, 5% CO2.

The transition time between resection in the operation room and arriving at the neurophysiology lab was less than 60 minutes. Immediately upon arrival at the lab, the tissue block was cleaned of residual blood and after identifying the pia-white matter (WM) axis, the pia was carefully removed. Subsequently, slices of 300 *µ*m-thick were made using a vibratome (Leica V1200S), in ice-cold aCSF solution. After recovering in a heat bath between 15-30 minutes at 34 °C, the slices were kept for 1 hour at room temperature before recording in aCSF, which contained (in mM): 125 NaCl; 2.5 KCl; 1.25 NaH2PO4; 1 MgCl2 6H2O; 2 CaCl2 2H2O; 25 NaHCO3; 25 C6H12O6, bubbled with carbogen gas (95% O2/5% CO2).

Borosilicate glass patch pipettes (Hilgenberg) were filled with intracellular solution, containing (in mM): 110 K-gluconate, 10 HEPES, 10 KCl, 4 Mg-ATP, 10 Na-Phosphocreatine, 0.3 GTP, and biocytin 3 mg/ml, pH adjusted to 7.2-7.3 with KOH, osmolarity 270-300 mOsm. Using this intracellular, the recorded cell was passively filled with biocytin during the patch clamp recording, allowing for subsequent staining of the cell.

### Topological analysis

Algebraic topology provides mathematical tools to characterize geometric shapes by encoding features that persist across length scales. The Topological Morphology Descriptor (TMD, (29)) represents the branching structure of trees as a persistence barcode, encoding the start and end distances from the soma of each branch in the underlying structure as an interval in the real line.

Given a rooted tree *T* with a set of *N* branches that consist of terminations or leaves *l* and intermediate branching points or bifurcations *b*, and given a real-valued function *f* on the nodes of the tree, such as the Euclidean (or path) distance from the root *R*, the TMD algorithm generates a persistence barcode *PB*:

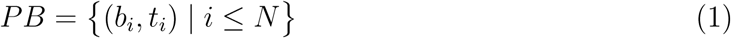

Each bar (*b_i_, t_i_*) in the persistence barcode associates a branch in the tree *T* with a pair of real numbers: if the *i^textth^* branch has bifurcation *b* and leaf *l*, then *b_i_* = *f* (*b*) and *t_i_* = *f* (*l*).

In brief, the persistence barcode is a set of intervals that encode the values of the function *f* at the start *b* and the end *l* of each branch. An equivalent representation of the persistence barcode is the corresponding persistence diagram *PD*, in which each interval of the barcode is encoded by the pair of its endpoints, seen as a point in the real plane.

There are many well-known methods to generate vectorizations of persistence barcodes, from which one can then compute various statistics. For example, the persistence diagram can be converted to a finite-dimensional vector representation, the persistence image *PI* (59), which is essentially a sum of Gaussian kernels centered around the points in the persistence diagram.

Another useful vectorization of a persistence diagram or barcode is its topological entropy (60), which is computed from the lengths of the bars:

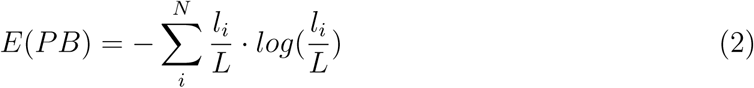

where *l_i_* = |*t_i_* − *b_i_*| is the length of each bar and *L* = Σ*^L^_i_ l_i_* is the total length of all bars.

### Bias analysis

Data from at least 115 different human patients were used in this study, but more than 500 human reconstructed neuronal morphologies.

The biological reconstructions of human pyramidal cells were analyzed to check for potential biases. Previous studies have shown that the morphology of human pyramidal cells correlates with intelligence and other cognitive traits (61; 9). Consequently, we examined whether any available metadata (sex, medical condition, age) correlated with the observed topological differences. This analysis is crucial to ensure that the differences between species are not artifacts stemming from the patients’ medical conditions or overlooked biases in the dataset. While sex and medical condition are binary variables in the metadata, age was categorized into two groups: below and above 40 years. Though age and cognition are expected to correlate with morphological characteristics, our goal here was to detect any potential bias in our results rather than to explore broader trends. Our analysis revealed no evidence of such biases.(Figure S2).

Additionally, we tested whether neurons from the same patients were more similar compared to those from different patients (Figure S3). Interestingly, our results showed that neurons from the same patients exhibited slightly less variability than those from different patients. However, the magnitude of this effect was not significant enough to influence our overall conclusions. Nevertheless, this finding presents an intriguing path for further investigation in the future.

### Topological scaling

The TMD associates a persistence diagram to any tree and thus to any reconstructed neuron morphology. Re-scaling a tree transforms the associated persistence diagram as follows. If *T* is a tree with a corresponding persistence barcode *PB* = (*b_i_, t_i_*) | *i* ≤ *N* given by the affine function *f*, the tree *T ^′^* is obtained from *T* by linear, or uniform, scaling by a factor *α >* 0 of all the branches, then its associated persistence barcode *PB^′^* is

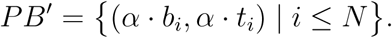

Note that the persistence entropy is scale-invariant as shown:

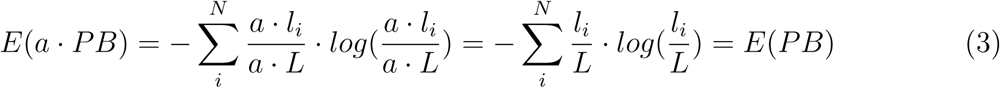

If *T* is a tree with a corresponding persistence barcode *PB* as above, and the tree *T ^′′^* is obtained from *T* by scaling by a pair of factors *α, β >* 0 of all the branches, so that the bifurcations of the branches are scaled by *α* and the leaves by *β*, then its associated persistence barcode *PB^′′^* is

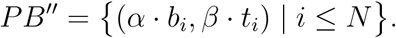

### Scaling optimization

To evaluate the scaling hypothesis, we have designed two experiments: the first one assumes uniform scaling through the morphology, and the second one non-uniform scaling as described above.

As a measure of the distance between two populations of cells, we computed the sum of the differences between the persistence images of the two populations, where the persistence image of a population is the sum of the persistence images of each cell in the population.

In the case of uniform scaling, we sought an optimal scaling factor *α >* 0 that transforms the population of mouse cells to human cells, minimizing the distance between the two populations.

For the non-uniform scaling, we sought two independent scaling factors *α >* 0 and *β >* 0 to transform the population of mouse cells to human cells, minimizing the distance between the two populations.

In both cases, a gradient descent approach was implemented to identify the optimal factors. The results are presented in Figure S7, along with the respective experimental data for comparison. Even though the identified optimal factors minimize the topological distances (difference between persistence images) between the two populations, they cannot capture the individual data points.

### Computation of inter-neuron distances

The inter-neuron distances were computed in two ways: by a mathematical formula and by a simulation of points generated according to the selected density and the respective spatial dimension. For the mathematical formula, we used equation (4) from (62).

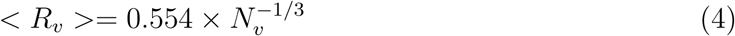

where 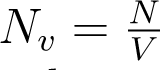 is defined as the number of particles *N* per unit volume *V*.

The result was confirmed by the computational generation of 100 instances of uniform point processes and computing the average minimum distance within each set of points. The reported results correspond to the distance computed by the simulation, but the computation using the mathematical formula (4) yielded the same values.

### Density of dendritic branches

Based on the total number of synapses within a cortical volume *S_v_* and the density of synapses *d_syn_* on the dendrites, the density *d_dend_* of dendritic branches within a cortical volume is computed as:

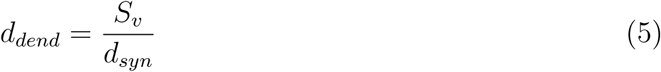

Similarly, the density of axons can be computed by the bouton density *d_bouton_*:

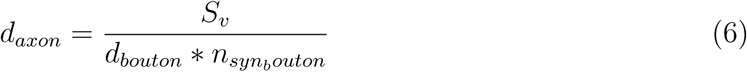

However, due to inconclusive experimental data about the number *n_synbouton_* of synapses that correspond to each bouton, we were unable to provide an accurate measurement for the respective axon density.

In addition, the measurements of axonal lengths and bouton densities for individual neurons are difficult to assess experimentally, due to the inability of staining techniques to accurately stain the axonal processes that are further away from the soma. Therefore, the analysis of axonal data should be re-assessed once more accurate experimental data become available.

### Memory capacity

Poirazi et al. (51) computed the memory capacity based on two different models: the linear model, which assumes dendritic inputs are summed linearly at the neuronal soma, and the non-linear model, which assumes the dendritic branches integrate the signal non-linearly. Independent of the choice of non-linearity introduced on the dendrites, memory capacity increases significantly if non-linearities are taken into account.

Given a dendrite with *m* branches, each branch contains *k* excitatory inputs, thus *s* = *mk* synaptic contacts and a set of *d* dendritic inputs, the memory capacity can be computed as follows.

For the linear model, the computed memory capacity is:

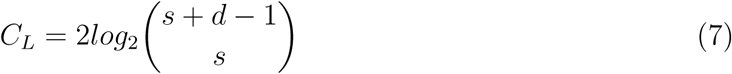

Note that in this case, the number of branches *m* and their synaptic length *k* are not relevant for the computation of the memory capacity.

For the non-linear model, the computed memory capacity is:

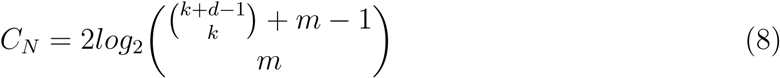

On the contrary, for the non-linear model, the distribution of synaptic lengths *k* on the *m* branches plays a crucial role in the memory capacity of the dendrite. As shown in Fig 14B the memory capacity increases with the number of branches, due to the increased number of ways to combine the dendritic inputs. The number of branches here corresponds to the number of computational subunits in the dendritic tree (63), rather than the physical branches of the tree.

### Morphological reconstructions processing

Morphological reconstructions are processed to correct common experimental errors, such as disconnected neurites from the soma, when the start of a neurite is not accurately recorded, zero diameters, when diameter information is missing, and similar small mistakes. In addition, pyramidal cells are aligned so that the apical dendrite is facing toward the positive *Y* −axis. These corrections result in a curated dataset of morphologies. For the generation of the circuit, morphologies are also unraveled to compensate for tissue shrinkage, and repaired for missing branches, to recover branches that were cut during tissue slicing. These corrections do not recover the original morphology, but better approximate the full extent of neuronal processes in the biological tissue.

An important disadvantage of morphological reconstructions in human tissue is the inability to recover fully intact axonal processes. As a result, most reconstructions are missing part of the axonal processes and some of them don’t have any axon reconstructed. For this reason, intact dendrites are matched with the best-reconstructed axons to recover as much as possible the missing axonal branches. We only choose to include in our dataset axons that have more than 20 branches to achieve more realistic connectivity in the circuits we build. Both dendrites and axons are then *cloned*, i.e. copied with a minimal added noise on their coordinates, to populate a cortical circuit of 10*K* morphologies from the small original dataset of 500 neurons. In the future, we intend to computationally synthesize (64; 65) neuronal morphologies instead of cloning them.

### Anatomical reconstruction of circuits

The circuits are generated as hexagonal columns with a predefined side length *α* and height *H*. To avoid boundary effects, seven columns are generated and the results are computed for the central column. For both species, the side length of the hexagon is 476*µm*. The height for the human columns is 1070*µm* and for the mouse is 235*µm*. For these dimensions, the respective cortical volume that will be generated is 0.6*mm*^3^ in humans and 0.13*mm*^3^ in mice. The somata are distributed homogeneously through the hexagonal column. According to the species details, cell densities are defined for inhibitory and excitatory cells. Since this simplified model aims to study the link between morphology and connectivity, specific density gradients are not considered. There is no distinction between layers 2 and 3 in densities.

For human circuits, the cell density is set to (≈ 25, 700*/mm*^3^) and (≈ 137, 600*/mm*^3^) for the mouse (11). Taking into account that the cell density for excitatory cells in the human cortex is 70%, we populate the human cortical circuit with 10500 pyramidal cells and 4500 interneurons. Respectively, taking into account that the cell density for excitatory cells in the mouse cortex is 85%, we populate the mouse cortical circuit with 15200 pyramidal cells and 2700 interneurons.

### Computation of appositions

The appositions between a pair of neurons were computed as the number of ”contact points” where the pre-synaptic axons approach the post-synaptic dendrites within a distance of 2*µm* (Figure S15). An apposition *A* is assigned to a pair of neurons *i, j* if and only if:

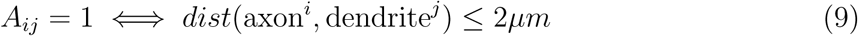

*A_ij_*= 1 ⇐⇒ *dist*(axon*^i^,* dendrite*^j^*) ≤ 2*µm* (9) It is important to note that these contact points do not directly represent synaptic connections, but rather indicate potential sites for synapse formation. Since appositions only represent potential synapses, they overestimate the actual synapses that are formed between neurons. The detailed methodology for the computation of the appositions is described in (27).

### Network processing

To bring the connection probability within a reasonable range, we applied a random reduction to the connectivity matrix formed by the appositions between neurons. To select the percentage of reductions, we computed the estimated connectivity density for both species for a network of 100K neurons. This adjustment ensured a more realistic representation of the network by reducing excessive connections. However, due to inconsistencies in the number of synapses per neuron in different experimental data (66; 28; 22) these values are indicative and should only be considered as a reference.

According to (22) the total number of synapses on a human pyramidal neuron is estimated to be 15K as opposed to the mouse neuron, which is 12K. From (30), we know that in the human the average number of synapses per connection is 4, while for the mouse it is 3.2. Interestingly this means that the average in-degree in both species is around 3, 750 cells:

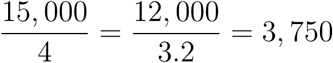

For 100K neurons the connection probability within the network is:

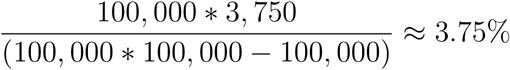

Therefore, we randomly reduce the networks of appositions, so that we have a biologically realistic connectivity network, around 4%. Note that in this paper, we do not perform any further reduction of appositions based on biological properties, as described in (27), due to the limitations of experimental data for human circuits for specific connection probabilities.

### Computation of simplex distribution

Recent advances in algebraic topology (26; 27) have introduced a powerful tool for comprehending network complexity - the computation of cliques and cavities formed by the underlying graph. This approach allows for a deeper understanding of the intricate connections and structures within networks, enabling researchers to unveil hidden patterns and gain insights into their organizational principles.

The dimension of a clique in a network is one less than the number of neurons of which it is composed; higher dimensions indicate higher complexity (Figure S1). In directed graphs, directed cliques are of particular significance as they comprise a single source neuron and a single sink neuron, representing a distinct pattern of connectivity, with the potential to influence the flow of activity within the network, making it an important consideration in network analysis (67; 38).

The mouse and human networks were partitioned into excitatory and inhibitory subnetworks. For each subnetwork, we computed the connection probability and analyzed the distribution of simplices, allowing us to assess the network characteristics and topological complexity specific to each subpopulation.

## Data availability

The data and the code used in this study is available in Zenodo 10.5281/zenodo.14258204. The topological analysis was performed using https://github.com/BlueBrain/TMD. The morphological analysis was performed using https://github.com/BlueBrain/NeuroM.

## Author contributions

L.K. conceived and supervised the study, designed the computational experiments, performed analysis, and wrote the manuscript. Y.S. supervised, performed, and validated reconstructed neurons. A.A. analyzed neuronal morphologies and generated neuronal networks. N.B. collected and revised literature information. R.B.P. reconstructed neurons and provided anatomical data for the human and mouse temporal cortex and hippocampus. J.C. edited the manuscript. J.D.F. provided anatomical data for the human and mouse temporal cortex. K.H. designed the topological methods and edited the manuscript. H.D.M. provided anatomical data for the human cortex. E.J.M. performed experiments and reconstructed human neurons. I.S. supervised the study. H.M. supervised the study and provided funding. C.d.K. provided data and supervised the study. J.M. contributed with histological preparations. R.P. and M.P. contributed with electrophysiological recordings of interneurons. R.T.D. and R.S. provided the human tissue for LNMC reconstructions. All authors reviewed the paper and discussed the results at all stages of this study.

## Acknowledgements

The authors thank Daniela Egas-Santader for her useful contributions to topological analysis of the networks. We also thank the visualization team of the Blue Brain Project for figure editing. This study was supported by funding to the Blue Brain Project, a research center of the École polytechnique fédérale de Lausanne (EPFL), from the Swiss government’s ETH Board of the Swiss Federal Institutes of Technology. H.D.M. and C.d.K. were supported by grant awards U01MH114812 (H.D.M.) and UM1MH130981-01 (H.D.M. and C.d.K.) from National Institute of Mental Health, grant no. 945539 (Human Brain Project SGA3, H.D.M.) from the European Union’s Horizon 2020 Framework Programme for Research and Innovation, NWO Gravitation program BRAINSCAPES: A Roadmap from Neurogenetics to Neurobiology (NWO: 024.004.012, H.D.M.), ERC AdG ‘fasthumanneuron’ 101093198 (H.D.M) and an NWO Open Competition grant (ENW-M2, project OCENW.M20.285, C.d.K.). R.S. was supported by Swiss National Science Foundation (IZLSZ3 148803, IZLIZ3 200297, IZLCZ0 206045, 31003A 138526) and Synapsis Foundation (2020-PI02). J.D.F. and R.B.P. were supported by Grant: PID2021-127924NB-I00 funded by Ministerio de Ciencia e Innovación / Agencia Estatal de Investigació (MCIN/AEI/10.13039/501100011033).

## Supplementary Information

### Dataset

- **INF**: Data from Vrije Universiteit Amsterdam, lab directed by Christiaan de Kock (12; 13).
- **LNMC**: Data from the Laboratory of Laboratory of Neural Microcircuitry, École Polytechnique Fédérale de Lausanne (EPFL), lab directed by Henry Markram (33).
- **CSIC**: Data from Laboratorio Cajal de Circuitos Corticales, Universidad Politécnica de Madrid and Instituto Cajal, lab directed by Javier DeFelipe and Ruth Benavides-Piccione (68; 41).
- **NMO:** Publicly available data, downloaded from neuromorpho.org.
- **AIBS:** Publicly available data, downloaded from Allen Institute for Brain Science (15).

**Figure S 1:**
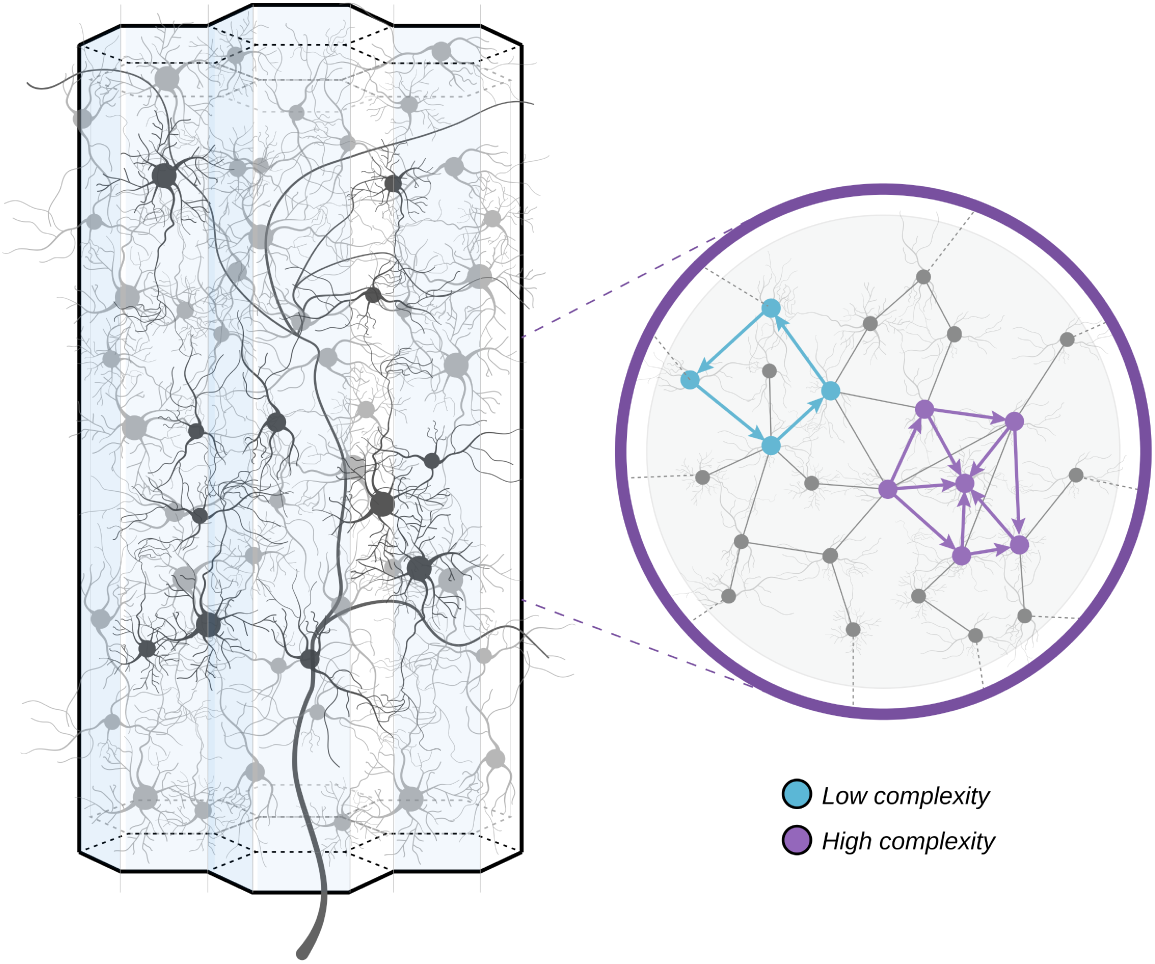
Illustration of topological complexity. Examples of low complexity (blue) for a cycle and high complexity (purple) for a 6-node simplex with one source and one sink.

**Figure S 2:**
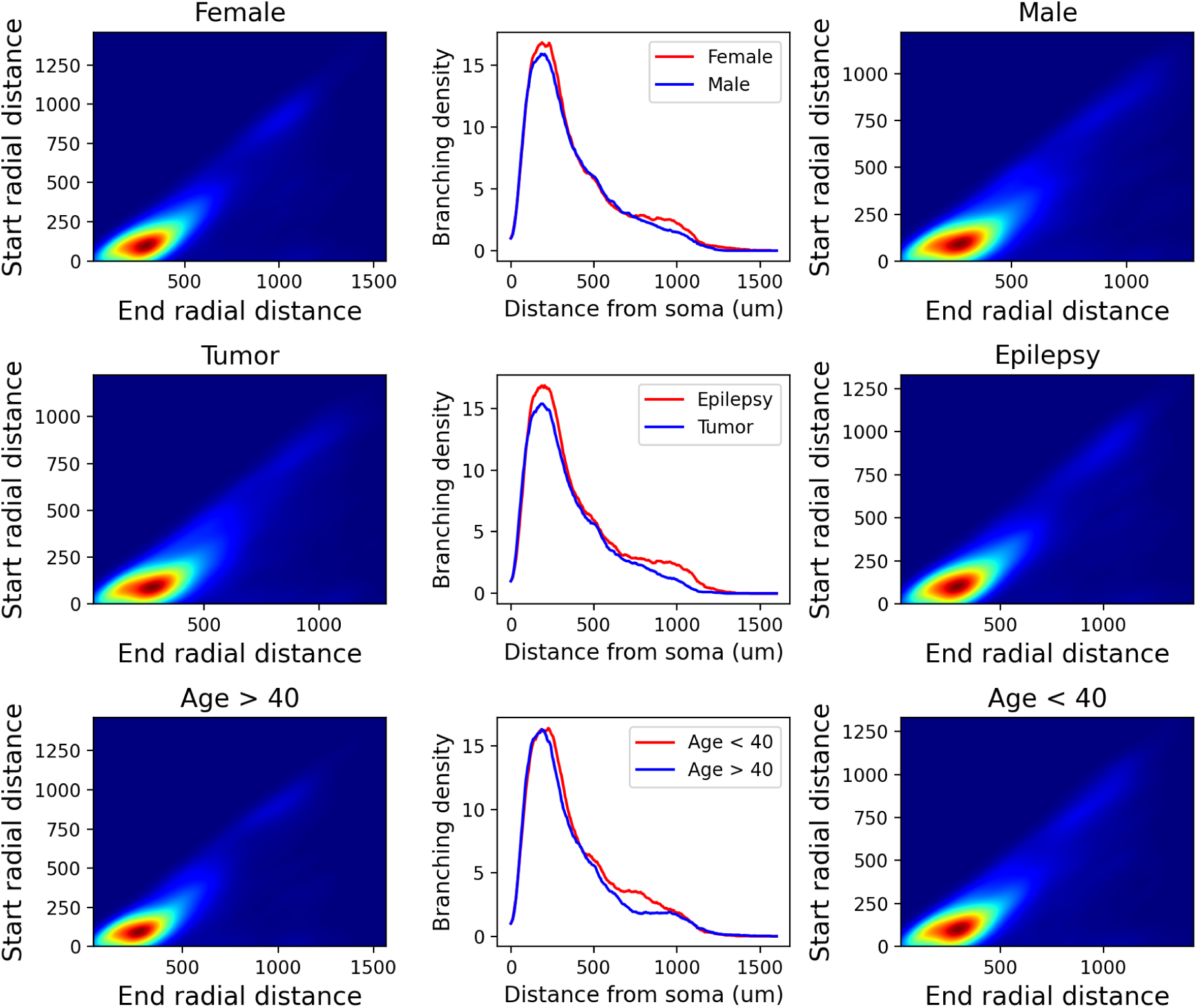
Analysis of morphological bias for sex, condition, and age. Persistence images of female patients are compared to male patients and their relative branching density (top). Persistence images of tumor patients are compared to epilepsy patients and their relative branching density (center). Persistence images for ages *<* 40 patients are compared to patients of *>* 40 and their relative branching density (bottom).

**Figure S 3:**
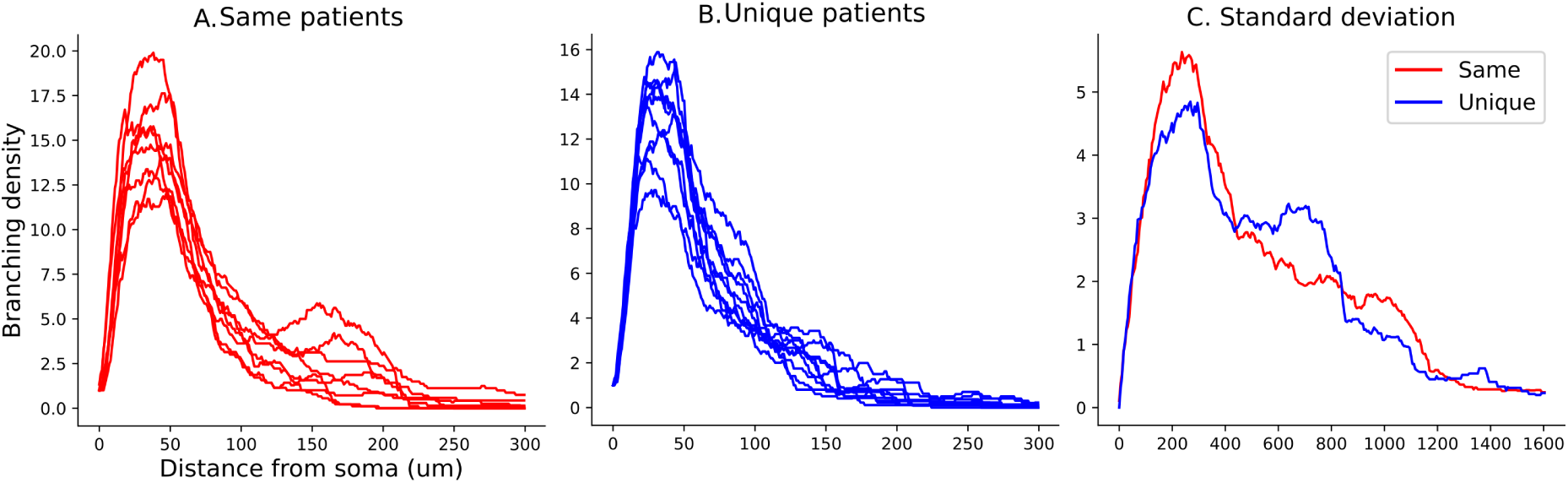
Intra versus inter-patient variance. The average branching density of neurons (A) from the same patients (7 − 10 neurons) is compared to the branching density of 10 randomly chosen neurons from unique patients (B). The standard deviation of branching density for neurons from the same patient is lower than the standard deviation from neurons of unique patients (C).

**Figure S 4:**
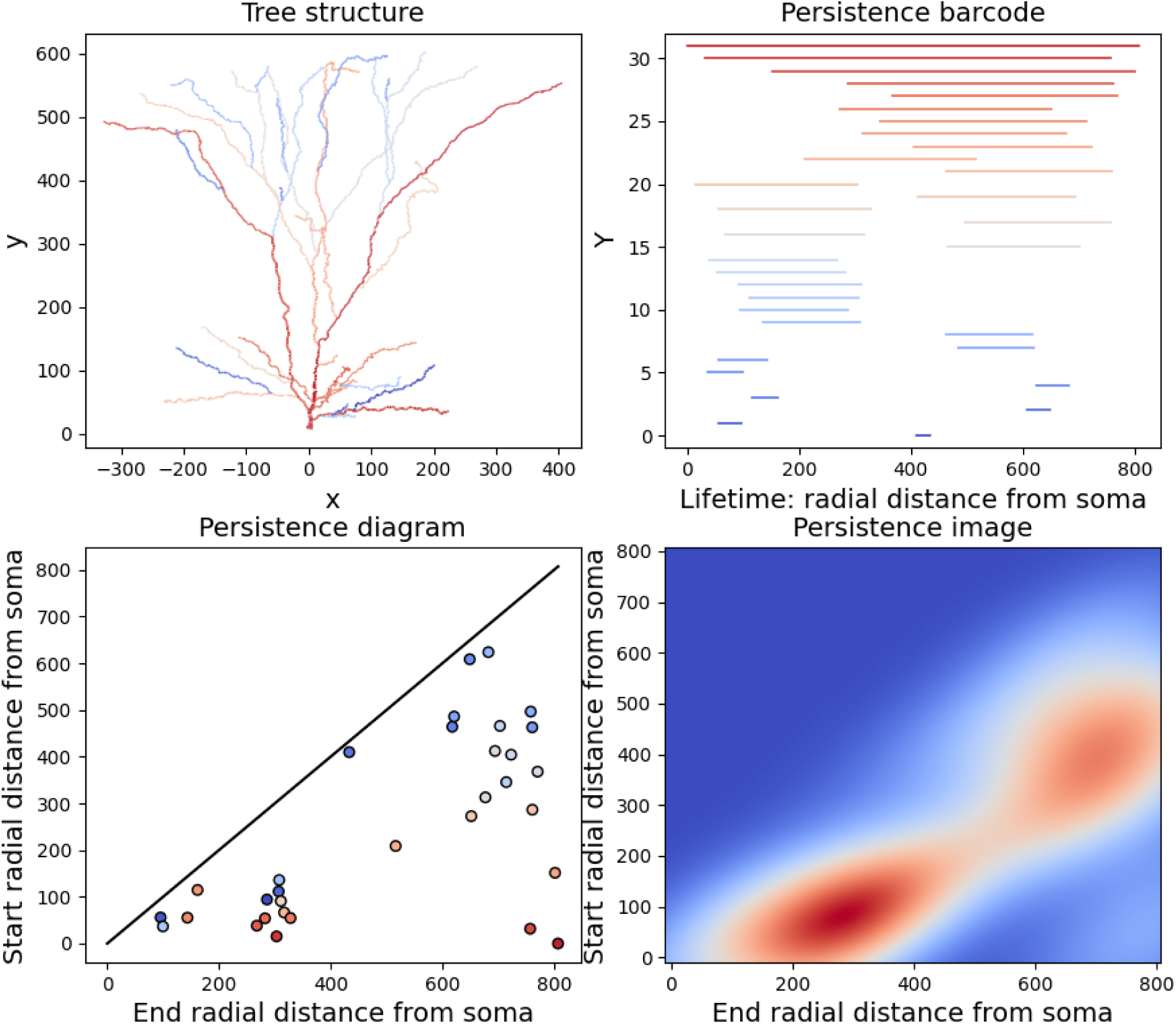
Topological morphology descriptor of an exemplar human layer 2 - 3 pyramidal cell apical dendrite. A. Apical dendrite, color-coded according to persistence components as illustrated in B. B. Persistence barcode, colormap from largest (red) to smallest branches (blue). C. Persistence diagram with the same color-code. D. Persistence image indicating areas of high density of branches (red) at different path distances from the soma (0,0).

**Figure S 5:**
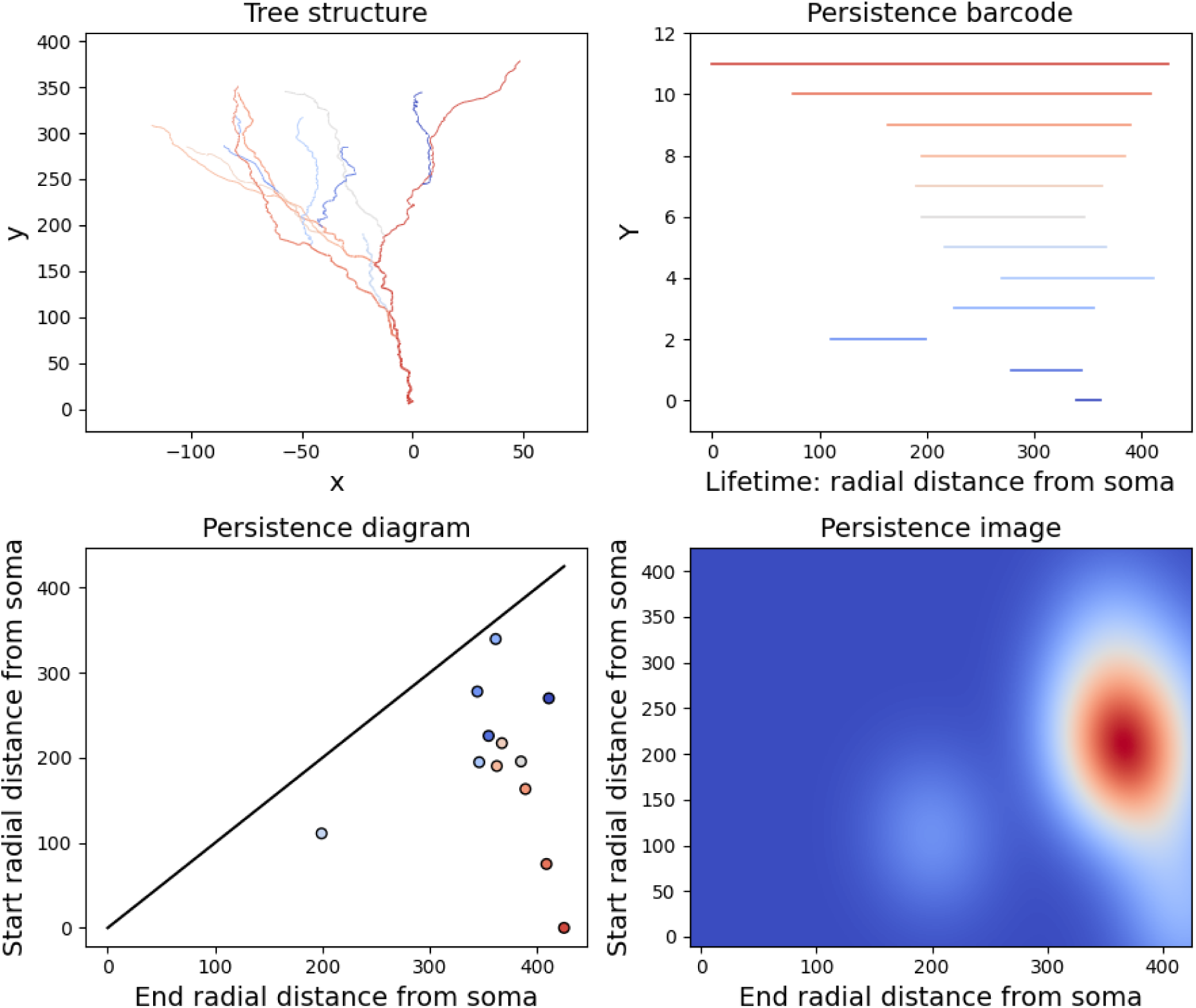
Topological morphology descriptor of an exemplar mouse layer 2 - 3 pyramidal cell apical dendrite. A. Apical dendrite, color-coded according to persistence components as illustrated in B. B. Persistence barcode, colormap from largest (red) to smallest branches (blue). C. Persistence diagram with the same color code. D. Persistence image indicating areas of high density of branches (red) at different path distances from the soma (0,0).

**Figure S 6:**
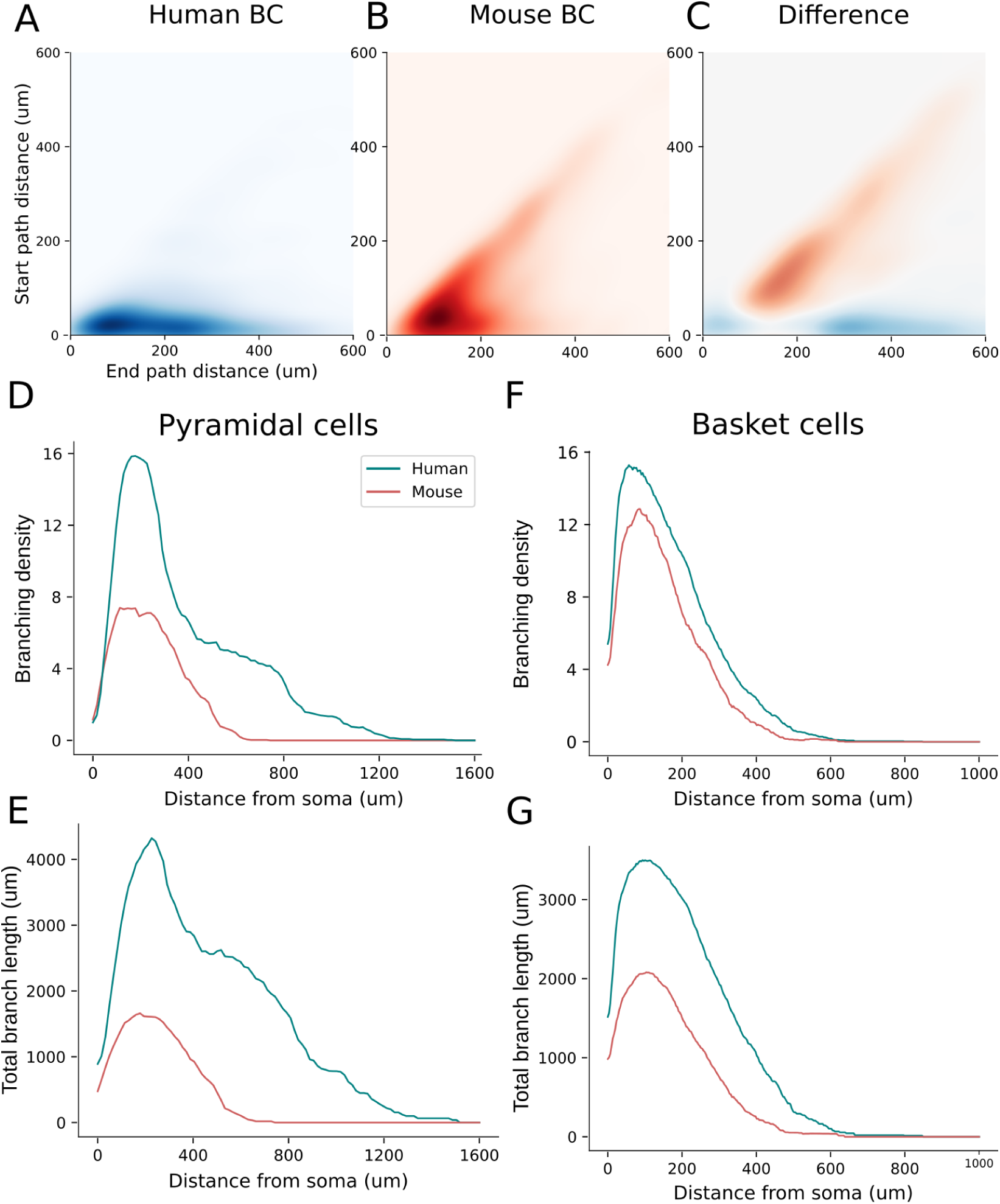
Topological analysis of mouse and human interneurons from cortical layers 2 and 3. A. Average persistence images for populations of human cells. B. Average persistence images for populations of mouse cells. C. shows the average difference between the persistence images of human (blue) and mouse (red) cells. The comparison of topological properties between pyramidal cells (D-E) and basket cells (F-G) of human (teal) and mouse (red) morphologies show that branching density is only marginally larger in basket cells (F), but branch lengths are significantly larger in human basket cells (G).

**Figure S 7:**
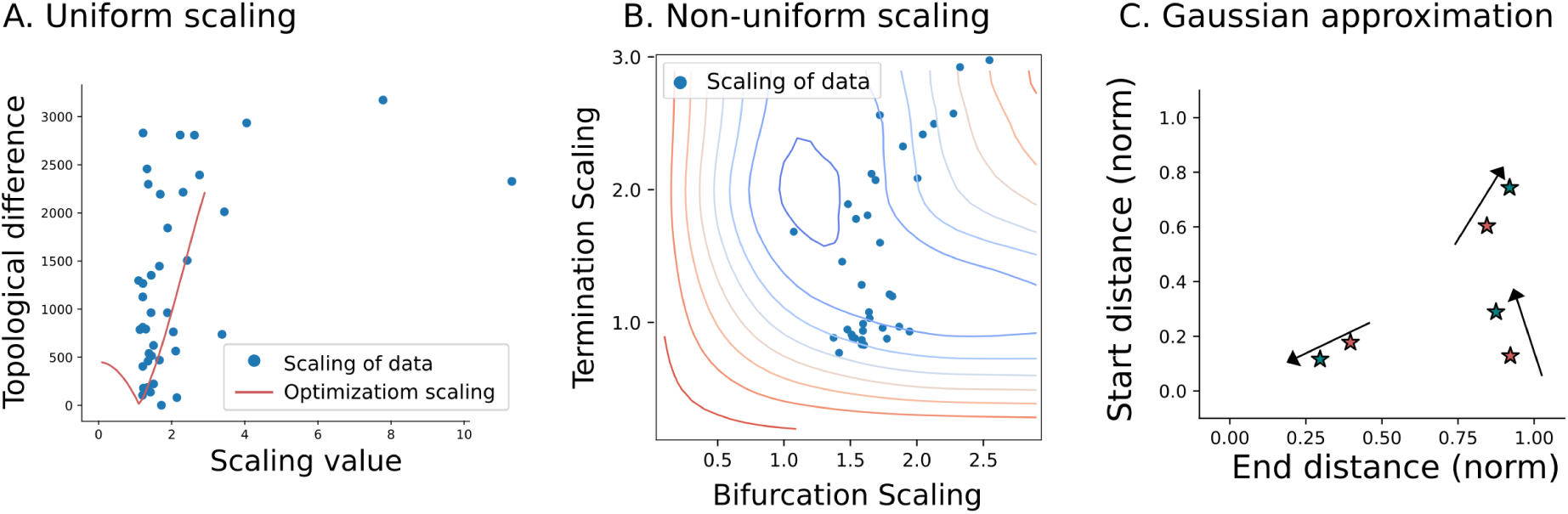
Optimization of scaling properties. A. Uniform scaling (red curve) compared to experimental data (blue points). B. Non-uniform scaling (contour) compared to experimental data (blue points). C. Persistence diagrams represent the start and end radial distances of branches from the soma. Gaussian kernels (3 centers) approximate the density of topological branching within the persistence diagrams of the two species (human: teal stars, mouse: red stars). The optimal transformations of the three Gaussian kernels to convert the normalized persistence diagrams from mouse to human are not aligned, indicating the absence of a consistence transformation between the species.

**Figure S 8:**
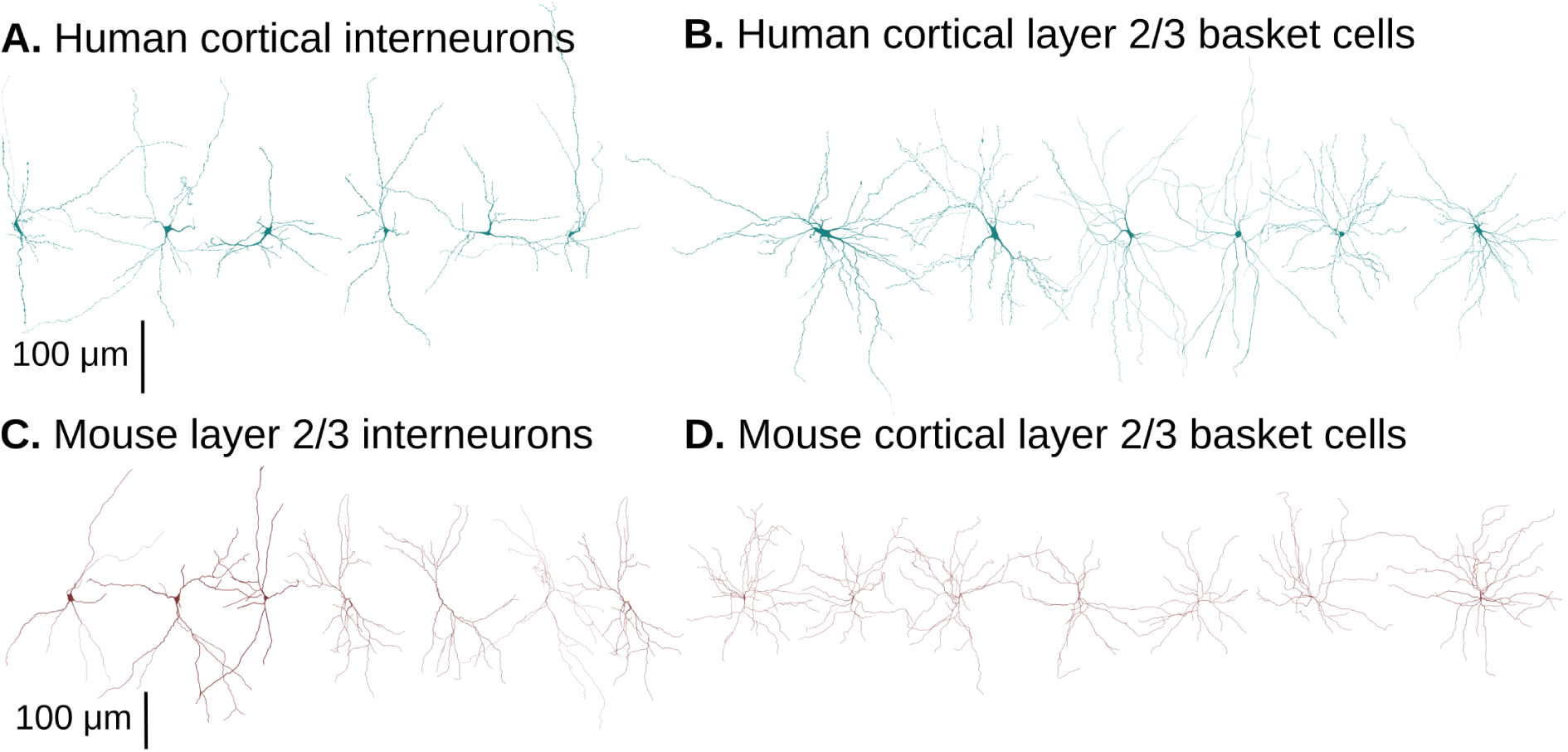
Examples of interneuron morphologies. A. Human cortical interneurons. B. Human layers 2 and 3 basket cells. C. Mouse cortical interneurons. D. Mouse layers 2 and 3 basket cells.

**Figure S 9:**
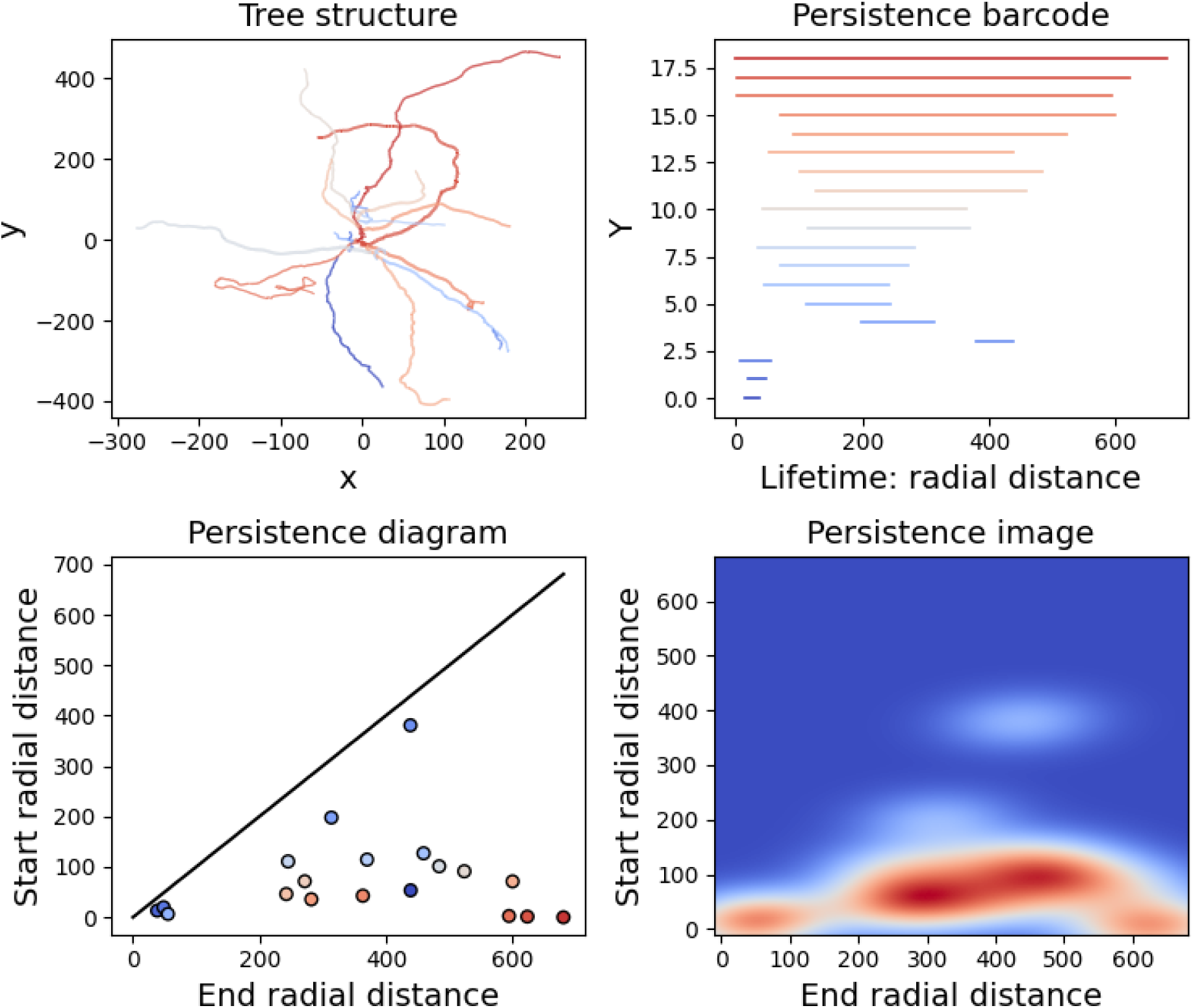
Topological morphology descriptor of an exemplar human cortical basket cell dendrites. A. Basal dendrite, color-coded according to persistence components as illustrated in B. B. Persistence barcode, colormap from largest (red) to smallest branches (blue). C. Persistence diagram with the same color-code. D. Persistence image indicating areas of high density of branches (red) at different path distances from the soma (0,0).

**Figure S 10:**
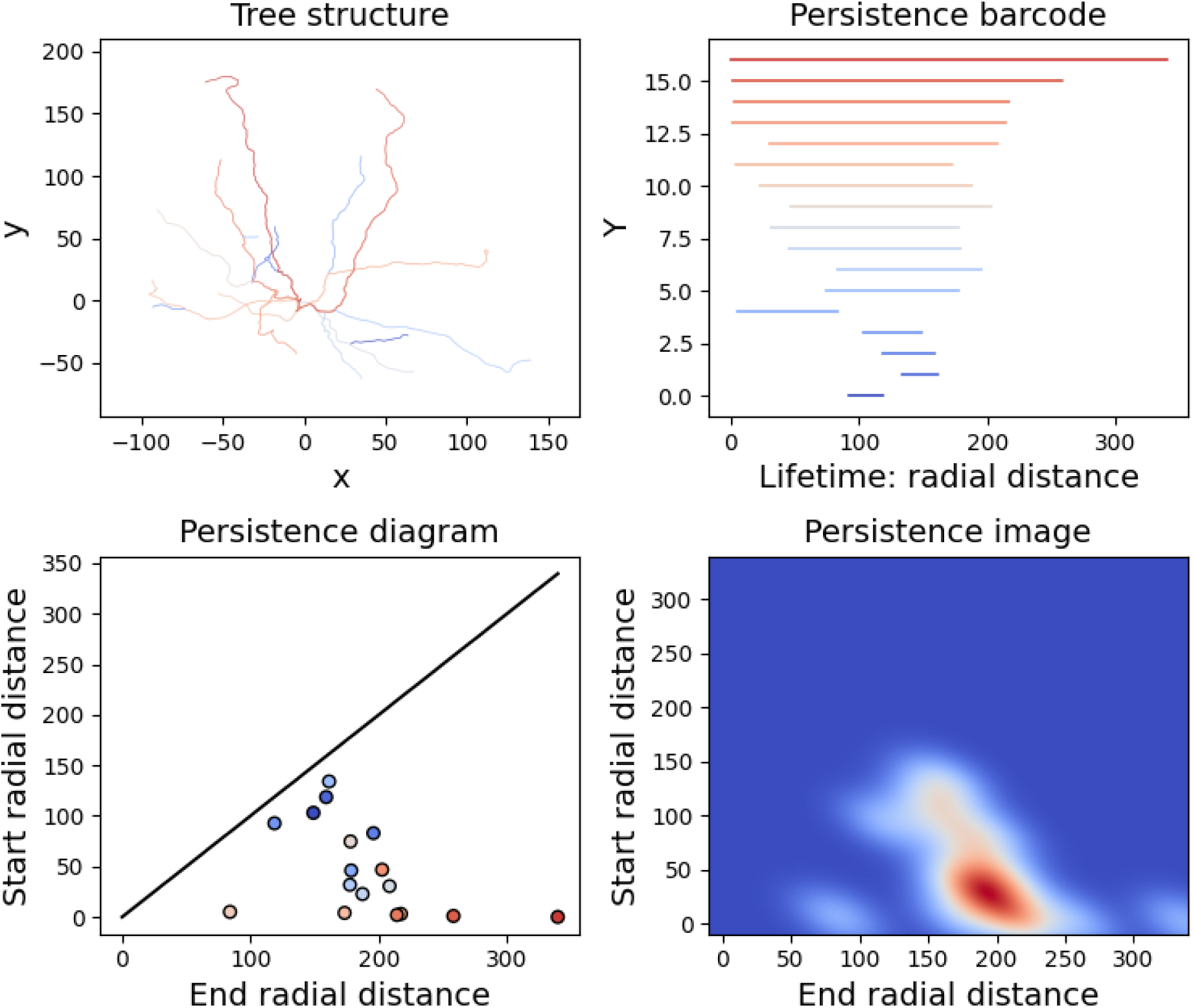
Topological morphology descriptor of an exemplar mouse cortical basket cell dendrites. A. Basal dendrite, color-coded according to persistence components as illustrated in B. B. Persistence barcode, colormap from largest (red) to smallest branches (blue). C. Persistence diagram with the same color code. D. Persistence image indicating areas of high density of branches (red) at different path distances from the soma (0,0).

**Figure S 11:**
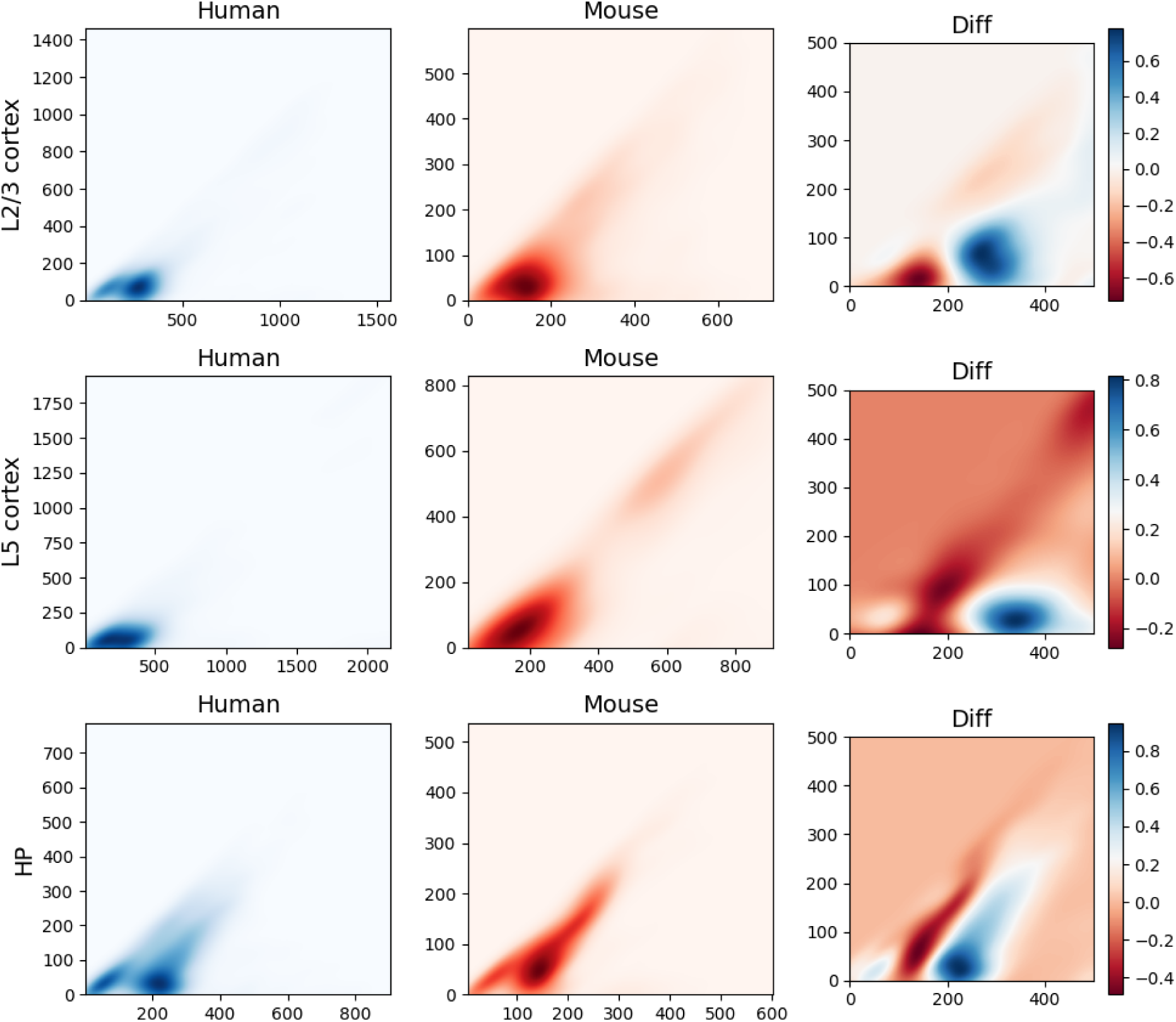
Topological analysis of mouse and human pyramidal cells from cortical layers 2, 3, and 5 and hippocampus. Column 1 shows the average persistence images for populations of human cells from different brain regions in blue. Column 2 shows the average persistence images for populations of mouse cells from different brain regions in red. Column 3 shows the average difference between the persistence images of human (blue) and mouse (red) cells from different brain regions. The topological differences that were observed between human and mouse pyramidal cells of layers 2 and 3 generalize to different layers and brain regions.

**Figure S 12:**
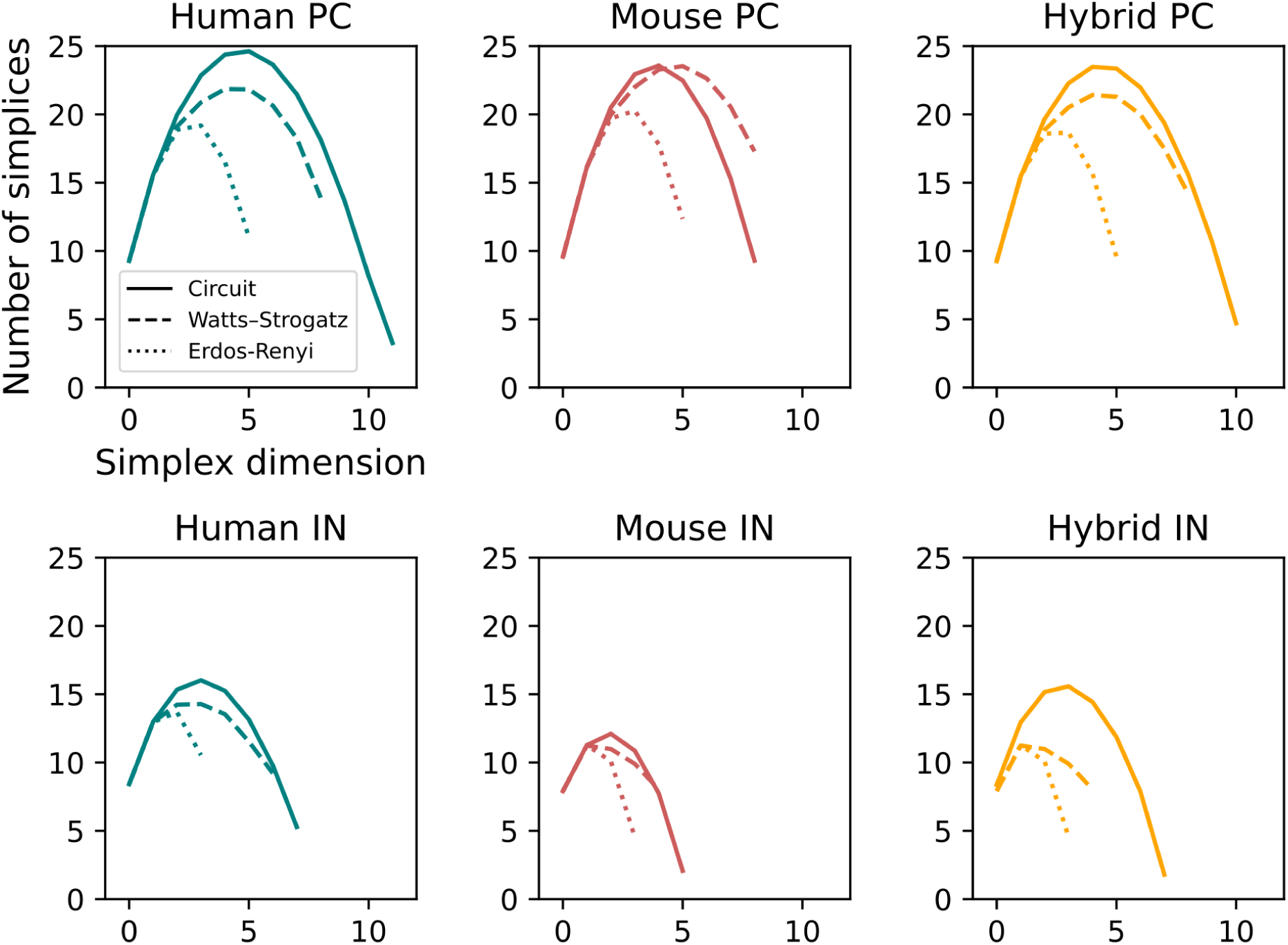
Comparison of simplice distribution to random networks. Simplices of human (pyramidal cells - PC and interneurons - IN), mouse (PC and IN) and hybrid (PC and IN) are compared to random networks based on the Watts-Strogatz and Erdos-Renyi models of the same network size and density.

**Figure S 13:**
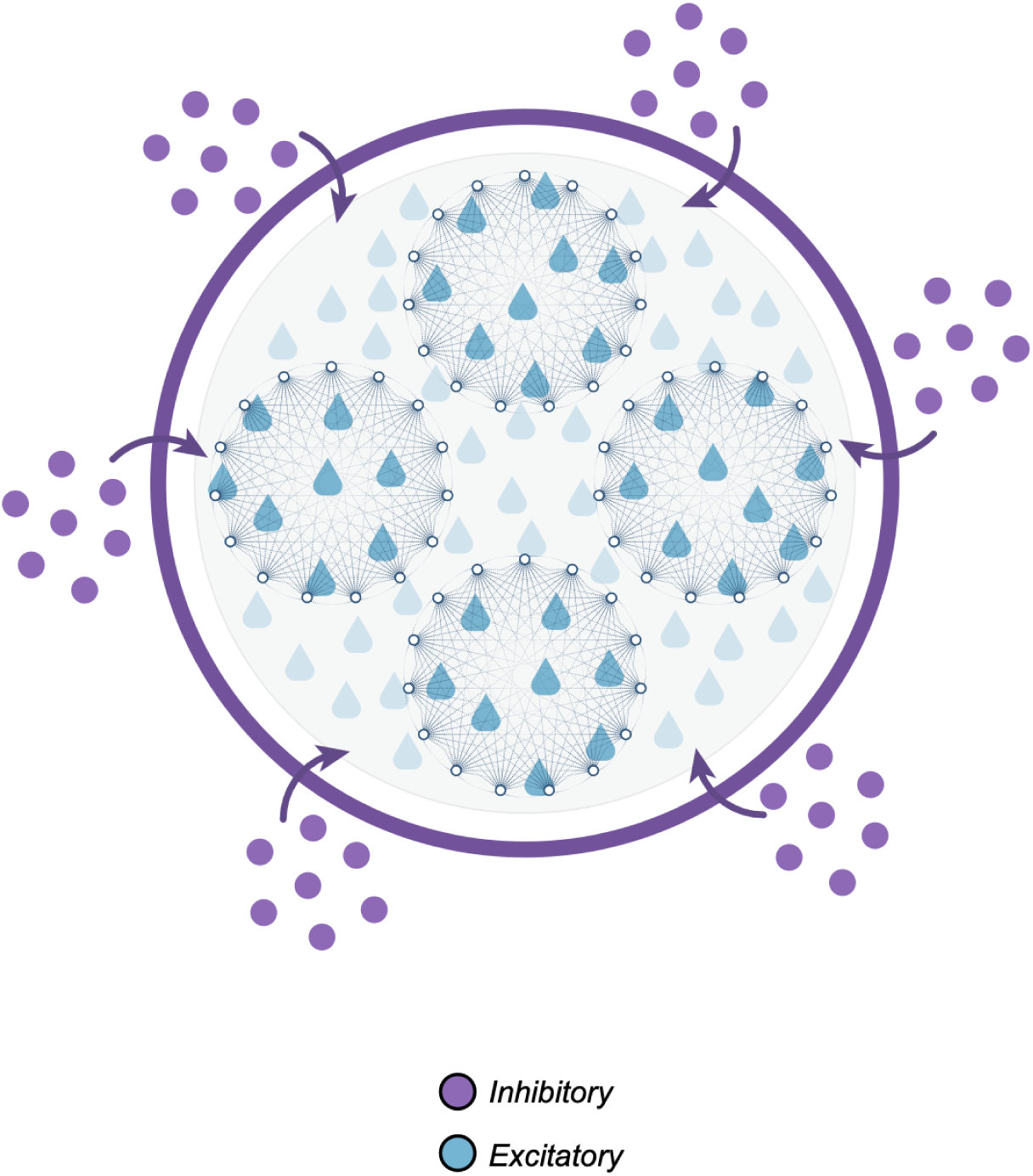
Schematic representation of interplay between interneurons and pyramidal cells. Strongly connected sub-networks of pyramidal cells (cliques) are controlled by small clusters of interneurons. More interneurons are needed to control the highly complex sub-networks of pyramidal cells and balance inhibition-excitation.

**Figure S 14:**
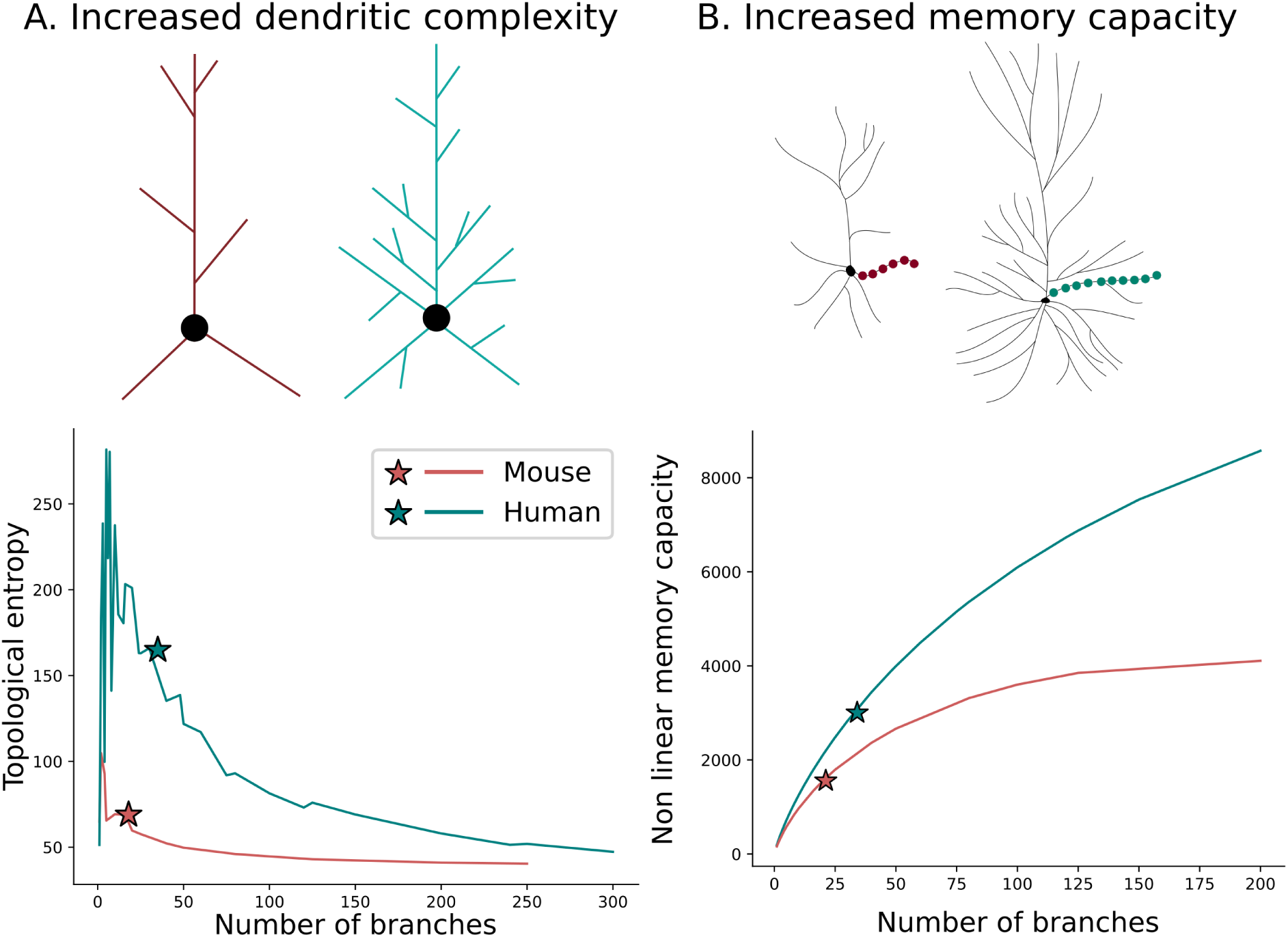
Memory capacity is enhanced by cell complexity. A. Cell complexity is measured by the topological entropy of the dendrites. The stars represent the average values for both species (red: mouse, teal: human). Topological entropy computed based on average number of branches is 1.8 times higher in human cells. B. Memory capacity is computed by the non-linear formula (see equation (9)) and depends on the number of branches and their lengths. The stars represent the average values for both species (red: mouse, teal: human). Memory capacity computed on average number of branches is 1.8 times higher in humans.

**Figure S 15:**
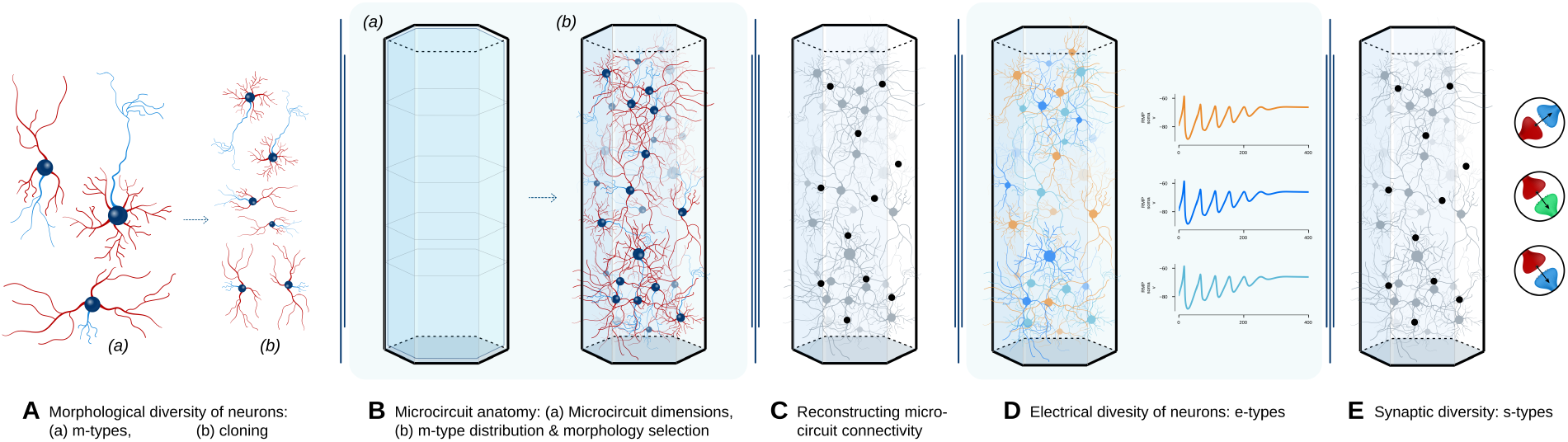
Complete pipeline for circuit generation. In this paper we implement the morphological and connectivity components without functional experiments. A. Morphological diversity. B. Microcircuit anatomy. C. Reconstructing connectivity. Electrical (D) and synaptic (E) diversity are not included in this paper.

**Table S 1:**
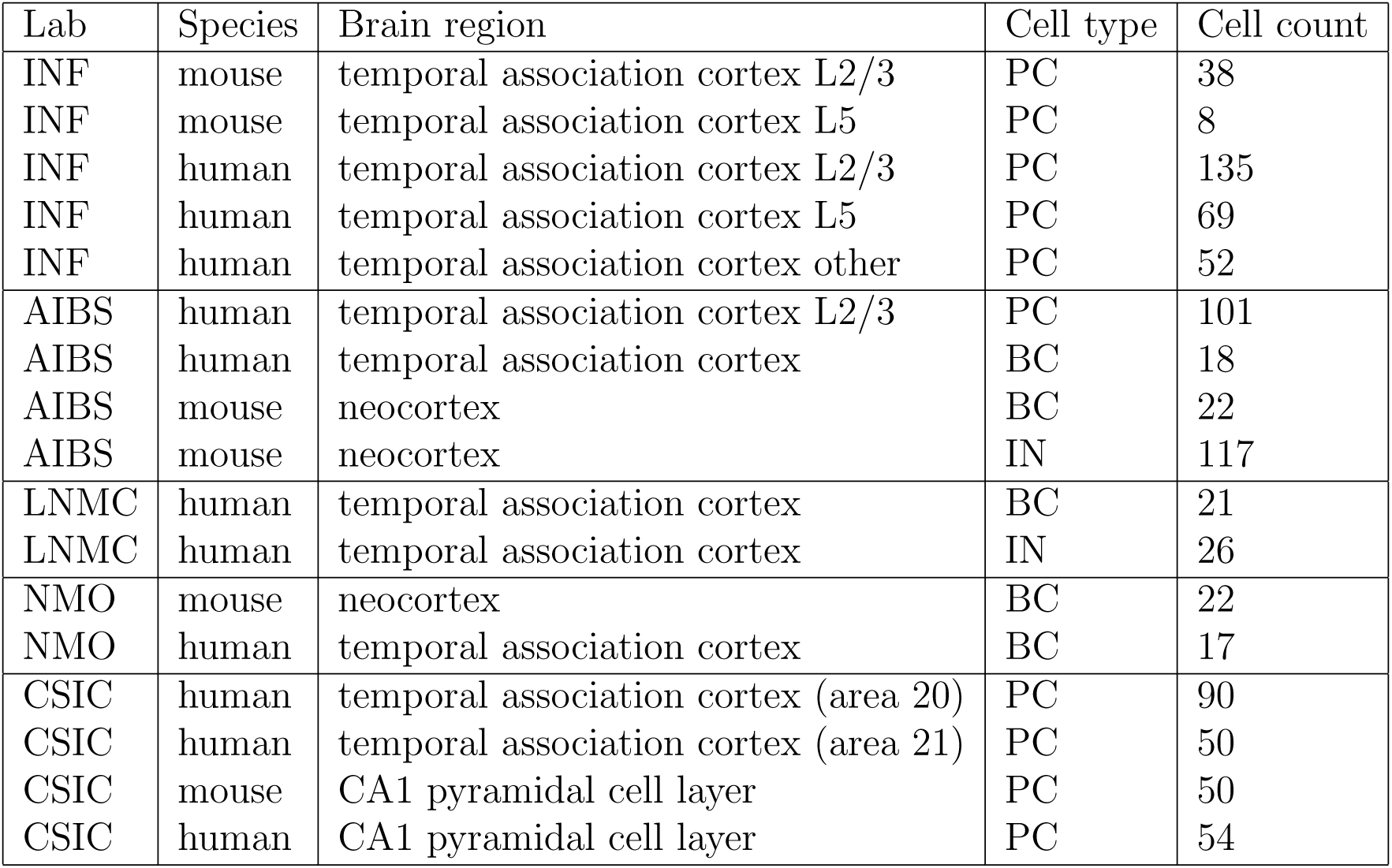
Summary of datasets of excitatory (pyramidal cells **PC**) and inhibitory cells (interneurons **IN**, basket cells **BC**).

| Lab | Species | Brain region | Cell type | Cell count |
| --- | --- | --- | --- | --- |
| INF | mouse | temporal association cortex L2/3 | PC | 38 |
| INF | mouse | temporal association cortex L5 | PC | 8 |
| INF | human | temporal association cortex L2/3 | PC | 135 |
| INF | human | temporal association cortex L5 | PC | 69 |
| INF | human | temporal association cortex other | PC | 52 |
| AIBS | human | temporal association cortex L2/3 | PC | 101 |
| AIBS | human | temporal association cortex | BC | 18 |
| AIBS | mouse | neocortex | BC | 22 |
| AIBS | mouse | neocortex | IN | 117 |
| LNMC | human | temporal association cortex | BC | 21 |
| LNMC | human | temporal association cortex | IN | 26 |
| NMO | mouse | neocortex | BC | 22 |
| NMO | human | temporal association cortex | BC | 17 |
| CSIC | human | temporal association cortex (area 20) | PC | 90 |
| CSIC | human | temporal association cortex (area 21) | PC | 50 |
| CSIC | mouse | CA1 pyramidal cell layer | PC | 50 |
| CSIC | human | CA1 pyramidal cell layer | PC | 54 |

**Table S 2:** Structural properties of networks for excitatory, inhibitory, and full networks. II: connections between inhibitory nodes, EE: connections between excitatory nodes for human and mouse.

| Networks | Human (II) | Human (EE) | Human | Mouse (II) | Mouse (EE) | Mouse |
| --- | --- | --- | --- | --- | --- | --- |
| Nodes | 4503 | 10621 | 15124 | 2724 | 14377 | 17101 |
| Edges | 431393 | 5717240 | 9661185 | 78766 | 10241482 | 11797379 |
| Density | 0.0213 | 0.0507 | 0.0422 | 0.0106 | 0.0496 | 0.0403 |
| In-degree (max) | 582 | 2070 | 3030 | 107 | 1377 | 1453 |
| In-degree (mean) | 95.80 | 538.30 | 638.80 | 28.92 | 712.35 | 689.86 |
| Out-degree (max) | 413 | 1600 | 2514 | 98 | 1343 | 1403 |
| Out-degree (mean) | 95.80 | 538.30 | 638.80 | 28.92 | 712.35 | 689.86 |
| Max dim | 8 | 12 | 12 | 6 | 9 | 9 |
| Peak dim | 3 | 5 | 5 | 2 | 4 | 4 |
| Simplices peak (log) | 16.02 | 24.62 | 25.55 | 12.10 | 23.58 | 23.67 |
| Simplices sum (log) | 16.75 | 25.49 | 26.40 | 12.67 | 24.23 | 24.33 |
| Complexity | 19.87 | 27.60 | 28.72 | 16.65 | 26.58 | 26.88 |

**Table S 3:** Average morphometrics for dendritic reconstructions of layers 2 and 3 for mouse (M) and human (H) pyramidal cells (PC) and basket cells (BC).

|  | HL23PC | HL23BC | ML23PC | ML23BC |
| --- | --- | --- | --- | --- |
| Num sections | 60.73 | 74.64 | 46.74 | 45.66 |
| Num bifurcations | 27.15 | 34.27 | 19.34 | 20.70 |
| Num leaves | 33.13 | 40.18 | 26.84 | 24.95 |
| Max radial distance | 301.25 | 383.63 | 208.09 | 264.15 |
| Total length | 5644.02 | 6431.81 | 2952.41 | 2756.53 |
| Section length | 91.06 | 102.09 | 62.88 | 62.35 |
| Mean radii | 0.32 | 0.32 | 0.34 | 0.31 |
| Section length (termination) | 143.72 | 154.02 | 89.40 | 82.35 |
| Section length (bifurcation) | 24.61 | 29.38 | 25.65 | 36.84 |
| Branch orders | 5.07 | 6.25 | 4.53 | 5.64 |
| Path distances | 141.57 | 168.20 | 102.52 | 154.12 |
| Radial distances | 109.94 | 124.07 | 79.62 | 120.99 |
| Section volume | 35.44 | 47.07 | 24.05 | 23.37 |
| Section areas | 183.12 | 215.30 | 133.69 | 122.12 |

**Table S 4:** Average morphometrics for apical reconstructions of layers 2 and 3 for mouse (M) and human (H) pyramidal cells (PC).

|  | HL23PC | ML23PC |
| --- | --- | --- |
| Num sections | 54.01 | 36.21 |
| Num bifurcations | 26.04 | 16.76 |
| Num leaves | 27.67 | 18.92 |
| Max radial distance | 692.36 | 349.13 |
| Total length | 5957.94 | 2523.44 |
| Section length | 109.24 | 70.66 |
| Mean radii | 0.34 | 0.38 |
| Section length (termination) | 160.03 | 93.67 |
| Section length (bifurcation) | 55.84 | 46.22 |
| Branch orders | 11.84 | 8.18 |
| Path distances | 291.68 | 227.77 |
| Radial distances | 240.83 | 185.69 |
| Section volume | 51.72 | 35.10 |
| Section areas | 236.73 | 167.18 |

**Table S 5:** Average morphometrics for axonal reconstructions of layers 2 and 3 for mouse (M) and human (H) pyramidal cells (PC) and basket cells (BC).

|  | HL23PC | HL23BC | ML23PC | ML23BC |
| --- | --- | --- | --- | --- |
| Num sections | 82.26 | 247.91 | 161.16 | 307.86 |
| Num bifurcations | 40.46 | 122.89 | 80.08 | 152.61 |
| Num leaves | 41.68 | 124.64 | 81.08 | 154.70 |
| Max radial distance | 1153.26 | 646.92 | 970.59 | 354.59 |
| Total length | 12090.53 | 17742.66 | 14906.03 | 8885.74 |
| Section length | 140.31 | 76.79 | 106.42 | 29.97 |
| Mean radii | 0.06 | 0.11 | 0.08 | 0.13 |
| Section length (termination) | 193.98 | 93.69 | 109.75 | 36.60 |
| Section length (bifurcation) | 80.21 | 59.42 | 102.84 | 23.31 |
| Branch orders | 11.84 | 18.29 | 13.29 | 18.86 |
| Path distances | 568.13 | 488.25 | 521.98 | 204.22 |
| Radial distances | 283.59 | 212.66 | 273.34 | 90.35 |
| Section volume | 4.5 | 4.34 | 2.13 | 1.82 |
| Section area | 59.4 | 56.73 | 51.98 | 23.97 |

**Table S 6:** Average morphometrics for neuronal reconstructions of layers 2 and 3 for mouse (M) and human (H) pyramidal cells (PC) and basket cells (BC).

|  | HL23PC | HL23BC | ML23PC | ML23BC |
| --- | --- | --- | --- | --- |
| Soma volume | 3184.75 | 2371.79 | 1522.11 | 1608.81 |
| Soma surface area | 868.05 | 814.38 | 630.33 | 619.7 |
| Soma radius | 7.71 | 7.85 | 7.02 | 6.77 |
| Max radial dist | 882.71 | 568.78 | 1073.61 | 409.26 |
| Mean width | 731.72 | 793.39 | 1189.56 | 391.73 |
| Mean height | 1148.49 | 838.69 | 1206.9 | 631.94 |

